# COP9 Signalosome Promotes Neointimal Hyperplasia via Deneddylation and CSN5-Mediated Nuclear Export

**DOI:** 10.1101/2023.04.11.536468

**Authors:** Samiksha Giri, Chao Suo, Ruggero Pardi, Gregory A. Fishbein, Khosrow Rezvani, Yabing Chen, Xuejun Wang

## Abstract

**BACKGROUND:** Neointimal hyperplasia (NH) is a common pathological response to vascular injury and mediated primarily by vascular smooth muscle cell (VSMC) migration and proliferation. The COP9 signalosome (CSN) is formed by 8 canonical subunits (CSN1 through CSN8) with its deneddylation activity residing in CSN5. Each or some of CSN subunits may have deneddylation-independent function. Despite strong evidence linking the CSN to cell cycle regulation in cancer cells, the role of the CSN in vascular biology remains obscure.

**METHODS:** Neointimal CSN5 expression in the lung tissue of pulmonary hypertension (PAH) patients was assessed with immunohistochemistry. Adult mice with smooth muscle cell-restricted CSN5 knockout (CSN5-SMKO) or CSN8 hypomorphism (CSN8-hypo) and cultured mouse VSMCs were studied to determine the role and governing mechanisms of the CSN in NH. NH was induced by ligation of the left common carotid artery (LCCA) and PDGF-BB stimulation was used to mimic the vascular injury in cell cultures.

**RESULTS:** Remarkably higher CSN5 levels were detected in the neointimal VSMCs of the pulmonary arteries of human PAH. LCCA ligation induced NH and significantly increased the mRNA and protein levels of CSN subunits in the LCCA wall of adult wild type mice. CSN5-SMKO impaired Cullin deneddylation and the nuclear export of p27 in vessel walls and markedly inhibited VSMC proliferation in mice. On the contrary, CSN8-hypo significantly exacerbated NH and VSMC proliferation *in vivo* and *in cellulo*.

Cytoplasmic CSN5 mini-complexes and the nuclear export of p27 were significantly increased in CSN8-hypo mouse vessels and cultured CSN8-hypo VSMCs. Nuclear export inhibition with leptomycin attenuated the PDGF-BB-induced increases in VSMC proliferation in both CSN8-hypo and control VSMCs. Further, genetically disabling CSN5 nuclear export but not disabling CSN5 deneddylase activity suppressed the hyperproliferation and restored p27 nuclear localization in CSN8 hypomorphic VSMCs. Interestingly, CSN deneddylase inhibition by CSN5i-3 did not alter the hyperproliferation of cultured CSN8-hypo VSMCs but suppressed wild type VSMC proliferation *in cellulo* and *in vivo* and blocked neointimal formation in wild type mice.

**CONCLUSION:** The CSN promotes VSMC proliferation and NH in injured vessels through deneddylation activity and CSN5-mediated nuclear export.

## Clinical Perspective

### What is new?

- A significant upregulation of CSN5 is discovered in the neointima VSMCs of human PAH and in mice after vascular injury and is required for VSMC proliferation and neointimal hyperplasia induced by vascular injury.
- Establishing both Cullin deneddylation and the nuclear export by CSN5 mini-complex as the underlying mechanisms for the promotion of VSMC proliferation and NH by CSN5.
- The first to demonstrate the promising prevention/suppressing effect of CSN5i-3, a small molecule inhibitor of the CSN, on injury-induced VSMC proliferation and NH.

### What are the clinical implications?

- CSN5 may be a novel therapeutic target to prevent or more effectively treat NH, a common pathological process underlying restenosis.
- Local application of a small molecular inhibitor of the CSN (e.g., CSN5i-3) to the vascular injury site should be explored to prevent restenosis and other maladaptive vascular remodeling where VSMC proliferation and NH are the primary culprit.

## INTRODUCTION

Cardiovascular disease (CVD) comprising of a range of disorders that detrimentally affect the heart and blood vessels, remains the leading cause of mortality and morbidity. A majority of CVD is caused by vascular dysfunction, either vascular proliferation or vascular rarefaction.^1–3^ Vascular smooth muscle cell (VSMC) proliferation is an important component of vascular remodeling and aberrant proliferation of VSMCs represents a major pathogenic factor of vascular diseases. Neointimal hyperplasia (NH) is a very common pathological process where the VSMCs migrate into the tunica intima layer and proliferate therein, thereby thickening the vessel wall and narrowing the vascular lumen.^4, 5^ This involves phenotypic switch of VSMCs, commonly occurs after vascular injury, and is considered the sole or major devastating post-operational complication for interventions like angioplasty,^5^ coronary artery bypass conduits^4^, stenting,^4, 6^ and other surgical repair.^7^ It also contributes to pulmonary hypertension (PAH),^1, 8^ diabetic vascular complications,^9^ and transplantation arteriopathy.^9, 10^ Therefore, it is extremely important to better understand the molecular mechanisms governing NH and thereby unveil new molecular targets for prevention and intervention.

The COP9 signalosome (CSN) is an evolutionarily conserved protein complex consisting of 8 canonical protein subunits (CSN1 through CSN8) that are encoded by the *COPS1* through *COPS8* genes. The *bona fide* function of the CSN is to regulate the Cullin-RING ligases (CRLs), the largest family of ubiquitin ligases, through a process known as deneddylation to remove NEDD8 from neddylated Cullin.^11–13^ The catalytic activity of the CSN is mediated by the zinc containing JAMM (JAB1-MPN-domain metalloenzyme) motif of CSN5/JAB1. However, CSN5 exerts its Cullin deneddylation function only when it is fully assembled in the CSN holocomplex consisting of all 8 canonical subunits.^12^ Interestingly, there is some evidence that CSN subunits can also exist in various forms of mini-complexes that are composed of some of the canonical CSN subunits and may have deneddylation-independent functions.^14, 15^ For example, a CSN5 containing mini-complex was purported to decrease nuclear levels of the cyclin-dependent kinase (CDK) inhibitor p27KIP1 (p27) by exerting its nuclear export in proliferating fibroblasts and Swiss 3T3 cells, independent of the deneddylation function.^16^

Considering its ability to deneddylate CRLs and thereby regulate the ubiquitin-proteasome system (UPS)-mediated degradation of cell cycle regulators, the CSN is a key regulator of cell cycle.^17^ Indeed, several experimental studies have shown great promise for a CSN inhibitor (CSN5i-3) to treat cancer.^18, 19^ CSN8 is the smallest and the least conserved CSN subunit.^13^ Decreasing CSN8 expression accelerated the cell growth rate in cultured mouse embryonic fibroblasts whereas cardiomyocyte-restricted ablation of CSN8 caused massive cardiomyocyte necrosis in mice.^20, 21^ Different from knocking down CSN8, silencing CSN5 showed severe growth inhibition effects on Hela cells, but myeloid-specific deletion of CSN5 exacerbated atherosclerotic lesion.^20, 22^ Regarding the pathophysiological significance of the CSN, most evidence gathered to date has been on the role of CSN5/JAB1 in cancer and highlighted a regulation by CSN5 on the degradation and subcellular distribution of tumor suppressors including p27 and p53.^12, 16, 23, 24^ Collectively, it appears that the CSN can either promote or inhibit cell cycle progression in a CSN subunit dependent and a cell-type specific manner. Regardless, the role of the CSN in vascular biology remains obscure and no reported study has examined the mechanistic contribution of the CSN to NH.

Here, we report that CSN5 is upregulated in VSMCs of human NH and experimental vascular injury triggers a marked upregulation of the CSN in VSMCs, which promotes VSMC proliferation and NH through both Cullin deneddylation and CSN5 mini-complex mediated nuclear export. On one hand, smooth muscle cell (SMC)-restricted CSN5 knockout (CSN5-SMKO) and the pharmacological inhibition of CSN deneddylase activity by CSN5i-3 suppressed NH and VSMC proliferation *in vivo* and *in cellulo*; on the other hand, CSN8 hypomorphism, which reduces moderately CSN holocomplex but increases CSN5 mini-complexes, exacerbated NH and VSMC proliferation in a manner that depends on CSN5-mediated nuclear export. Thus, the present study unveils the CSN as a major regulator of vascular remodeling after injury and identifies a novel strategy to intervene NH.

## METHODS

A full description of methods can be found in the online supplement.

### Animals

*Cops5* encodes CSN5. The creation of mice harboring Cops5-floxed alleles was described.^25^ The Myh11-CreERT2 transgenic mouse was obtained from The Jackson Laboratory [strain # 019079; Tg (Myh11-icre/ERT2)1Soff/J].^26^ To achieve homozygous smooth muscle-restricted CSN5 knockout (CSN5-SMKO), adult *Cops5^flox/flox^::Myh11-CreERT2* mice with littermate *Myh11-CreERT2* mice as the control (CTL) were injected intraperitoneally with tamoxifen (1mg/day x 5 days/round x 2 rounds, 2-day break in between rounds). Since the *Myh11-CreERT2* transgene is inserted into the Y chromosome, only male mice were suitable for studies involving CSN5-SMKO. CSN5-SMKO mice in the C57BL/6J in-bred background were used.

Generation of the CSN8 hypomorphic allele has been described.^20^ The neomycin gene inserted in an intron of a CSN8-floxed allele (CSN8^neoflox/+^) reduces CSN8 gene expression but this reduction of CSN8 gene expression does not cause CSN8 protein reduction until the other CSN8 allele is deleted (CSN8^neoflox/-^); hence, the CSN8^neoflox/-^ mice have been confirmed as CSN8 hypomorphic and the littermate CSN8^neoflox/+^ mice were used as controls. CSN8-hypo mice in the FVB/N inbred background were used.

All protocols for the care and use of animals in this study were approved by the IACUC of the University of South Dakota and in accordance with the NIH guidelines for the care and use of laboratory animals.

### Ligation of the left common carotid artery (LCCA)

To produce NH *in vivo*, LCCA ligation was performed on adult mice as described.^27^ The right common carotid artery (RCCA) was used as the uninjured intra-animal control vessel.

### Immunohistochemical assessment of CSN5 in human pulmonary arteries

Human lung tissue samples were taken from archived surgical pathology paraffin embedded tissue acquired as standard of care. The control tissue represents sections of normal segment of lung tissue taken during surgical resection of lung tumors. The PAH tissue was acquired from explanted lungs from patients with idiopathic PAH undergoing transplantation. These paraffin-embedded tissue sections were used for CSN5 immunohistochemistry (IHC) with the Universal 1-Step Polymer-Based IHC/DAB kit (Catalog# VB-6023D, VitroVivo Biotech, Rockville, MD).

### Cell cultures

VSMCs were enzymatically isolated from abdominal aortas of adult mice as described.^28^ VSMCs at passage 3-6 were used for all *in cellulo* experiments. Before each experiment, the cells were serum-starved and then stimulated with platelet-derived growth factor-BB (PDGF-BB, 10ng/ml).

### Statistical analysis

Data were statistically analyzed using GraphPad Prism (Version 9, GraphPad Software). All quantitative data are presented as mean±SEM unless otherwise indicated. Differences between the two groups were evaluated using two-tailed unpaired Student’s *t*-test or Mann-Whitney test whichever appropriate, based on normality tests. Difference among three or more groups were evaluated using one-way or, when appropriate, two-way ANOVA followed by Tukey’s pairwise comparison tests as indicated in figure legends. The *p* value or adjusted *p* value <0.05 is considered statistically significant.

## RESULTS

### 1. Upregulated CSN expression in vascular walls and VSMCs by injury

To better understand the significance of the CSN in vascular biology, we assessed CSN expression in the wall of LCCA proximal to the ligation site in wild type (WT) mice 1 week after LCCA ligation. The steady-state protein levels of all 6 representative CSN subunits were significantly higher in the ligated side (LCCA) compared with the counter non-ligated side (RCCA) (**Figure 1A, 1B**). Significant increases in the mRNA levels of all examined CSN subunits (CSN5, CSN6 and CSN8) were observed in the ligated side as well (**Figures 1C**, **S1**). Immunoprobing for CSN5, CSN6, and CSN8 in native proteins fractionated by native-PAGE, a technique separating native protein complexes based on their size, consistently revealed remarkable increases in the CSN holocomplex in the ligated LCCA (**Figure 1D**). Moreover, immunofluorescence confocal microscopy detected that the upregulated CSN5 (**Figure 1F**) and CSN8 (**Figure S2**) were primarily localized in the neointimal VSMCs. Taken together, these data indicate that the expression of CSN subunits and the CSN holocomplex are significantly upregulated in the SMCs of the neointima induced by vessel injury.

**Figure 1.**
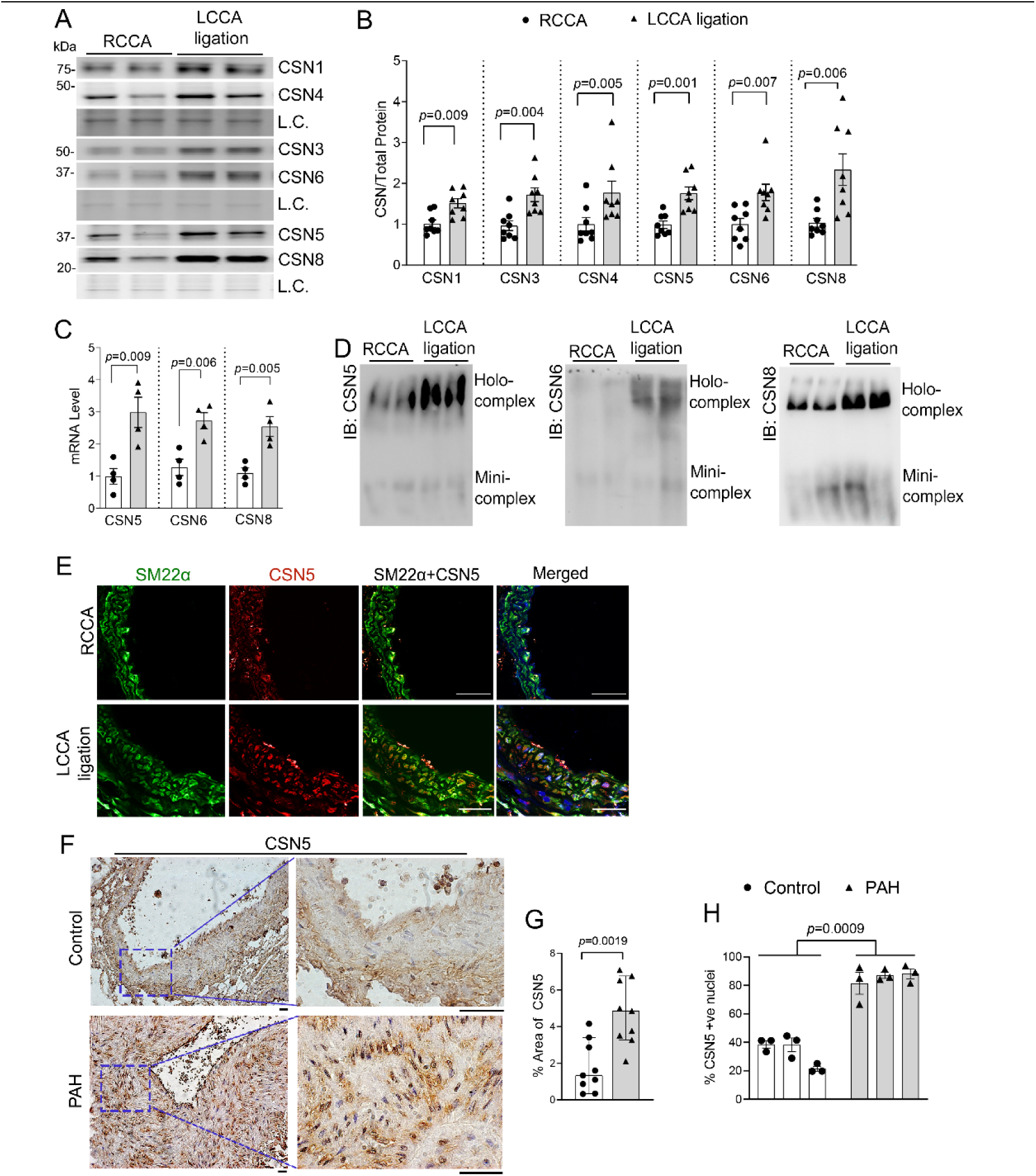
CSN subunits are upregulated in wild type mice after ligation of the left common carotid artery (LCCA) and in neointima of human pulmonary arterial hypertension (PAH) lesions. **A∼E**, Adult wildtype mice were subject to LCCA ligation; 1 week later, the LCCA proximal to the ligation site as well as the corresponding segment of the right common carotid artery (RCCA) were collected. **A** and **B**, Representative images (A) and pooled densitometry data (B) of western blot analyses for the indicated CSN subunits. L.C., a segment of the in-lane loading control image derived from the stain-free total protein imaging technology; this is the same for other figures. **C**, Real time PCR (qPCR) data of the indicated genes. Scatter plots superimposed by mean±SEM; each dot represents a mouse, Student’s t-test. **D**, Representative images of immunoblotting (IB) for the indicated CSN subunits fractionated by native PAGE. **E**, Representative confocal micrographs of immunofluorescence staining for CSN5 (red) and SM22α (green). Nuclei were stained with DAPI (blue). Scale bar= 75μm. F∼H, Paraffin sections of lung tissues from humans with PAH or without (Control) were processed and immunohistochemically stained for CSN5 (brown). **F**, Representative images of CSN5 immunohistochemistry. **G and H**, The percent CSN5-positive area (G) and the percentage of CSN5-positive nuclei (H) of smooth muscle cells in the tunica media (Control) or the neointima (PAH) of the pulmonary arteries. Scale bar=100μm. Scatter plots superimposed by Median±95%CI (G) or Mean±SEM (H); each dot represents a pulmonary artery, 3 arteries/patient, and 3 patients/group are included. *P* values are derived from Mann-Whitney test (G) or nested *t*-test (H).

Next, we were able to validate the CSN upregulation in human NH. The demographic information of the patients is shown in **Supplemental Table II**. Immunohistochemistry (IHC) revealed that CSN5, the enzymatic subunit of the CSN, was remarkably higher in the neointimal VSMCs of the pulmonary arteries in the explanted lungs of humans with PAH than in the VSMCs of control pulmonary arteries (**Figures 1F**, **S3**). Quantitative image analyses showed that both the percent of CSN5-positive areas and the percent CSN5-positive VSMC nuclei in the neointima of PAH tissues were significantly greater than those in the tunica media of control pulmonary arteries (**Figure 1G, 1H**).

### 2. Suppressing NH formation and VSMC proliferation by CSN5-SMKO in mice

We successfully achieved tamoxifen-induced CSN5-SMKO in adult *Cops5^fl/fl^::Myh11-CreERT2* mice (**Figures 2A**, **S4**). Consistent with loss of Cullin deneddylation activity, only the neddylated forms of Cul1 and Cul2 were discernible whereas Skp2 was substantially decreased in the vessel walls of both LCCA (ligated) and RCCA in CSN5-SMKO mice compared with littermate CTL mice (**Figure 2A**). Skp2 is an F-box protein that can serve as the substrate receptor in the Skp1-Cul1-Fbox complex (SCF^skp2^) to target p27 for ubiquitination and subsequent degradation by 26S proteasomes.^29^ The decrease of Skp2 was accompanied by a tendency of increased p27 (**Figure 2A, 2B**).

**Figure 2.**
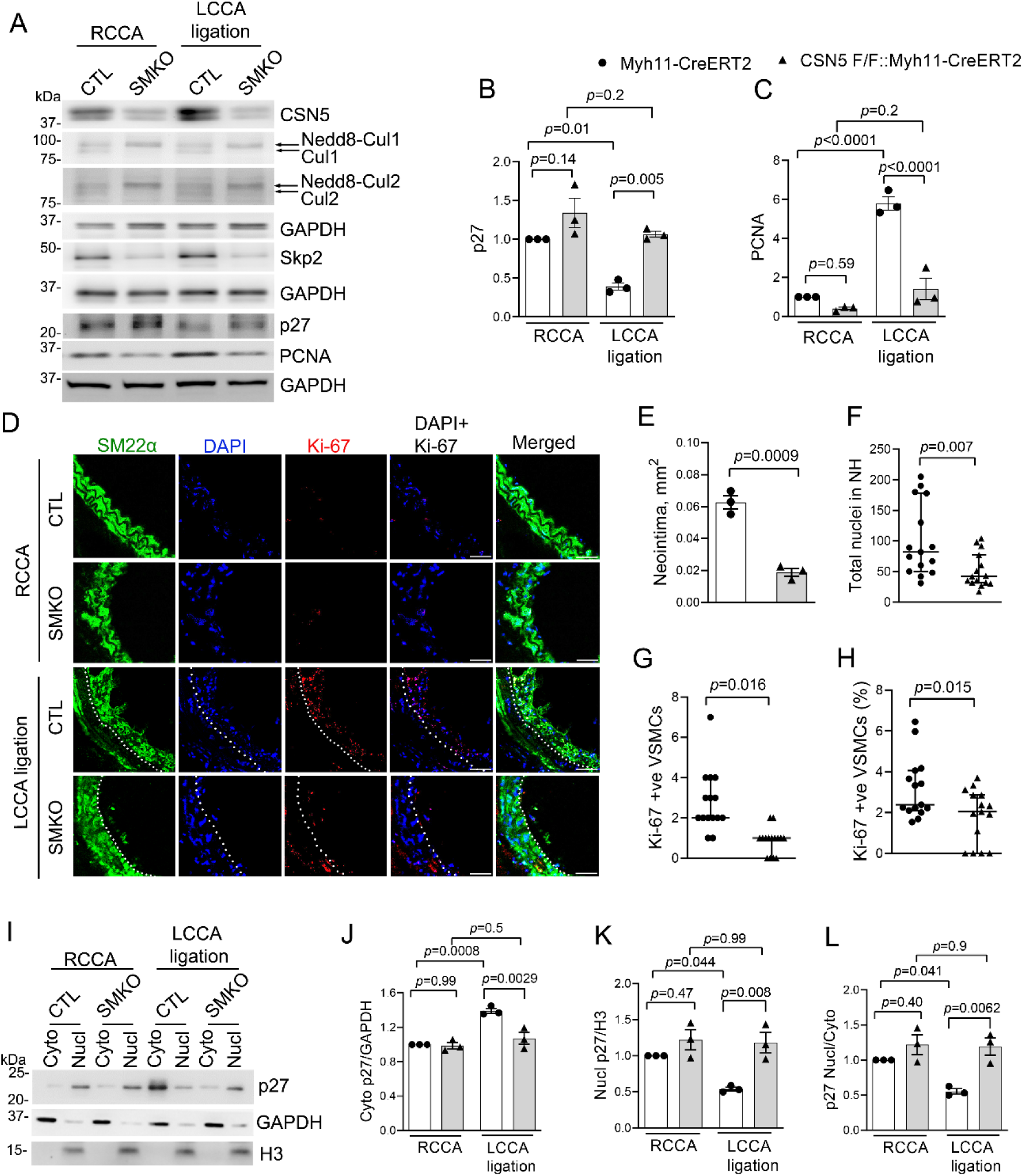
Smooth-muscle cell restricted CSN5 knockout (SMKO) suppresses VSMC proliferation. Homozygous SMKO and Myh11-CreERT2 transgenic (CTL) mice were subject to LCCA ligation; the LCCA proximal to the ligation site and the corresponding segment of unligated RCCA were collected 1 week later. **A∼C**, Representative images of western blot analyses for the indicated proteins (A) and pooled densitometry data for p27 (B) and PCNA (C). **D∼H**, Cross-sections of carotid arteries were immunofluorescence stained for SM22α (green) and Ki-67 (red) and counter-stained with DAPI to identify nuclei (blue). Shown are representative confocal images (D) and quantitative analysis (E-H) of neointima formed (E), total nuclei in neointima (F), Ki-67 positive VSMC in neointima (G), and the percentage of Ki-67 positive VSMC (H) calculated by normalizing the Ki-67 positive VSMC/total nuclei in the neointima. Scale bar=50μm. Scatter plots superimposed by Median±95%CI; each dot represents a section, 5 sections/mouse, 3 mice/group; Mann-Whitney test (F∼H). **I**∼**L**, Represenative images (I) and pooled densitomtery data (J-L) of western blot analyses for p27 in the nuclear (Nucl) and cytosolic (Cyto) fractions. RCCA was used as a baseline control. GAPDH and Histone H3 (H3) are probed as the loading control for cytosolic and nuclear proteins, respectively. Scatter plots superimposed by Mean±SEM; each dot represents a mouse; n= 3 mice/group (B, C, E, J-L); two-way ANOVA followed by Tukey’s tests (B, C, J-L) or unpaired Student’s *t*-test (E).

To study the necessity of CSN5 for NH, we characterized the CSN5-SMKO and CTL mice one week after LCCA ligation. Corresponding to the increased p27, the ligation-induced increase in the cell proliferation marker PCNA was significantly less in the CSN5-SMKO mice than that in the CTL mice (**Figure 2A**-**2C**). Double-immunofluorescence staining for cell proliferation marker Ki-67 and SMC marker SM22α revealed a significantly thinner neointima, fewer nuclei in neointima, fewer Ki-67-positive VSMCs per section, and lower percentage of Ki-67-positive nuclei in the neointima of CSN5-SMKO mice, compared with those in the CTL mice (**Figure 2D-2H**).

Further, we determined if ablation of CSN5 would affect the subcellular localization of p27 after vascular injury. p27 in the nucleus inhibits CDKs, but as the cell enters the proliferative phenotype, nuclear p27 gets translocated to the cytoplasm.^30^ Western blot analyses of nuclear fractions of mouse arteries revealed that LCCA ligation led to a significantly reduced nuclear p27 protein level in the CTL mice, but this was not observed in CSN5-SMKO mice (**Figure 2I-2L**). Taken together, these experiments using CSN5-SMKO mice demonstrate that CSN5 in SMCs is required for the reduction of nuclear p27, the elevation of VSMC proliferation, and the formation of neointima in response to vascular injury. No significant difference in VSMC proliferation was observed between CSN5-SMKO and CTL mice at baseline.

### 3. Exacerbated NH and VSMC proliferation in response to vascular injury in CSN8 hypomorphic mice

Since CSN5 harbors the deneddylase site for the CSN, the effect of CSN5-SMKO on NH could result from loss of the Cullin deneddylation activity, of CSN5-containing mini-complex, or of both. To decipher these possibilities, we turned to the CSN8 hypomorphic (CSN8-hypo) mice where all CSN subunits are moderately downregulated but the CSN5 mini-complex is not.^31^ Therefore, we investigated the effect of CSN8 hypomorphism on NH. Four weeks after LCCA ligation, histopathological examination showed LCCA neointimal thickening was markedly greater in CSN8-hypo mice than in CTL mice (**Figure 3A – 3E**) and this differential response was comparable between male and female mice (**Figure S5**). One-week after LCCA ligation, PCNA protein levels in the LCCA of CSN8-hypo mice were more than doubled compared to the CTL mice although no difference was detected from the RCCA that was not ligated (**Figure 3F, 3G**), indicating that CSN8 hypomorphism promotes vascular cell proliferation in response to vessel injury. Immunofluorescence confocal microscopy confirmed that LCCA ligation induced a markedly greater increase in Ki-67 positive VSMCs (**Figure S6**, **3H-3K**) and thicker neointima (**Figure 3H**, **S7**) in the LCCA of CSN8-hypo mice compared with those in the CTL mice. Between CSN8-hypo and CTL groups, no difference in PCNA proteins was observed in the RCCA of LCCA-ligated mice (**Figure 3F, 3G**) or the LCCA and RCCA of mice subjected to LCCA sham surgery (**Figure S8**). These results indicate that NH and VSMC proliferation are increased in CSN8-hypo mice after vascular injury.

**Figure 3.**
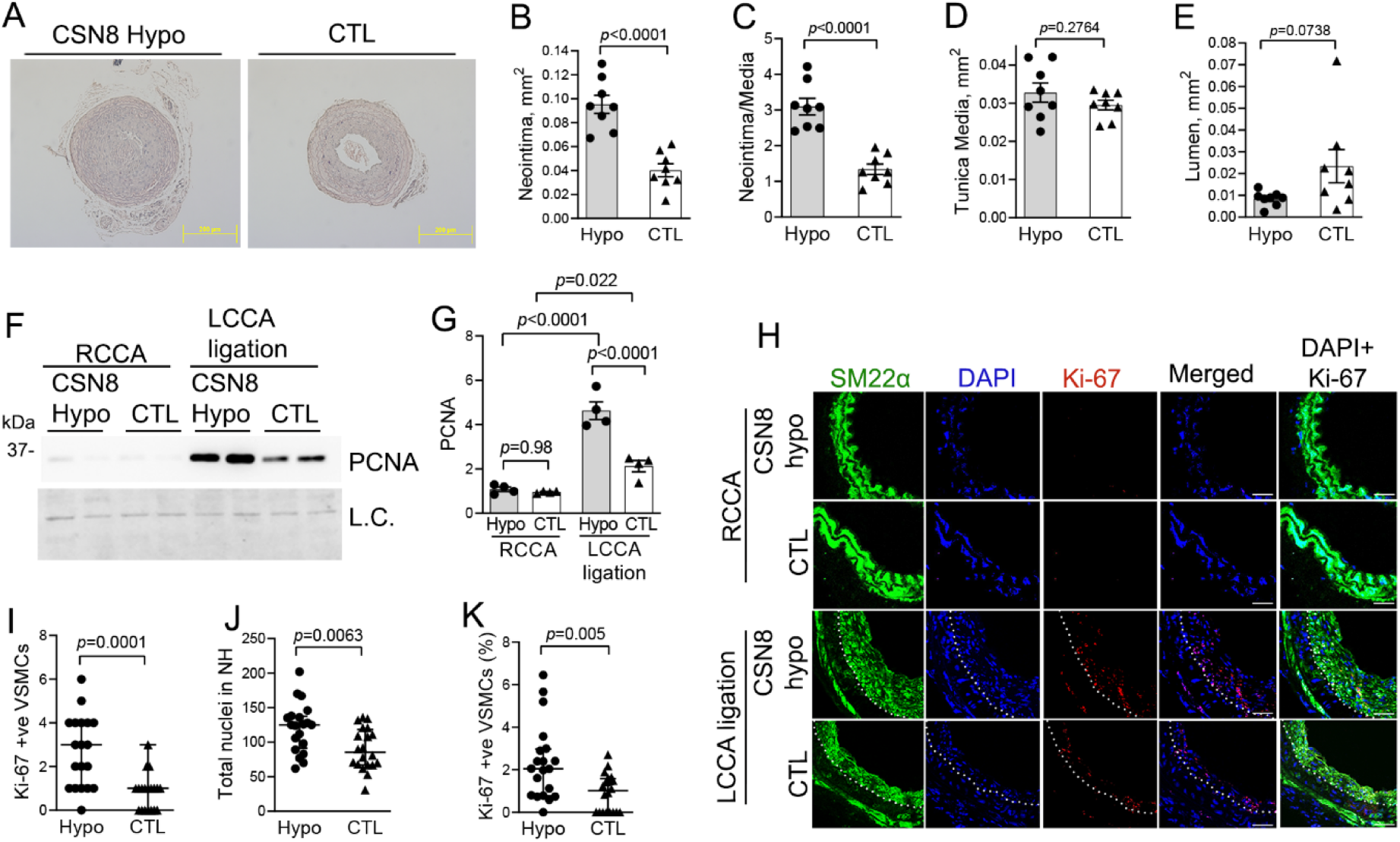
CSN8 hypomorphism (CSN8-Hypo) exacerbates LCCA ligation-induced neointimal thickening and vascular smooth muscle cell (VSMC) proliferation. CSN8-hypo and control (CTL) mice were subject to LCCA ligation; LCCA (ligated) and RCCA were collected 4 weeks (A∼E) or 1 week (F∼K) after ligation. **A∼E**, Representative images (A) and morphometric analyses of the indicated parameters (B∼E) from the H&E-stained cross-sections of paraffin-embedded LCCAs. Scale bar=200µm; scatter plots superimposed by mean±SEM; each dot = a mouse, n= 8 mice/group (4 males, 4 females); two-tail unpaired students *t*-test. **F** and **G**, Representative image (F) and pooled densitometry data (G) of western blot analyses for PCNA. Stain-free total protein imaging was used for loading control (L.C.). Scatter plots superimposed by mean±SEM; n= 4 mice/group; two-way ANOVA followed by Tukey’s tests. **H**, Representative confocal images of SM22α (green) and Ki-67 (red) immunofluorescence. SM22α was stained (green) to identify VSMCs and nuclei are counter-stained with DAPI (blue). Scale bar=50µm. **I∼K**, Quantitative data of Ki-67 positive VSMCs (I) and the total number of nuclei (J), as well as the percentage of Ki-67 positive VSMC (K) in the neointima of LCCAs. Scatter plots superimposed by median±95%CI; each dot represents a section; 5 sections/mouse; n=4 mice/group, Mann-Whitney test.

### 4. Decreased nuclear p27 and increased CSN5 minicomplexes in CSN8 hypomorphic vessels

To investigate if CSN8 hypomorphism affects the subcellular localization of p27, nuclear and cytoplasmic fractions were extracted from the carotid artery segments of the CSN8-hypo and CTL mice 1 week after LCCA ligation. In both LCCA and RCCA, nuclear p27 proteins were significantly lower, cytoplasmic p27 proteins were higher, and the nuclear p27 to cytoplasmic p27 ratios were significantly smaller in CSN8-hypo mice compared to CTLs (**Figure 4A, 4B**). These findings are consistent with the increased VSMC proliferation induced by LCCA ligation in CSN8-hypo mice (**Figure 3**).

**Figure 4.**
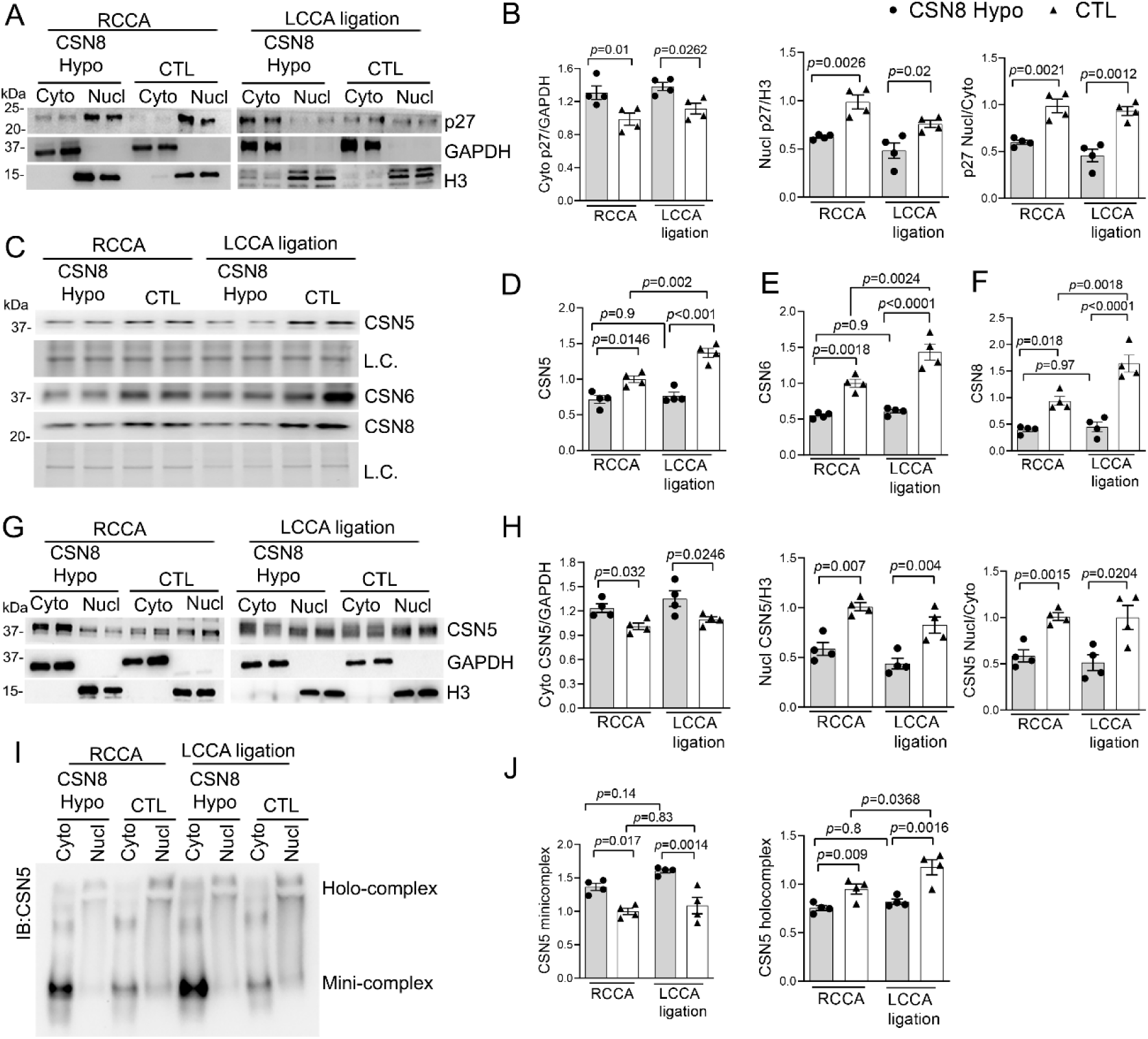
p27 and CSN5 is increased in the cytoplasmic fraction of CSN8-Hypo mice. LCCA and RCCA were collected from CSN8-hypo and control mice 1 week after LCCA ligation. For nuclear fractionation, three carotid arteries of the same group were combined into one sample. **A** and **B,** Representative images (A) and pooled densitometry data (B) of western blot analyses for cytoplasmic (Cyto) and nuclear (Nucl) p27. GAPDH and Histone H3 were probed as the loading control for cytoplasmic and nuclear fractions, respectively. **C∼F**, Representative images (C) and pooled densitometry data of the western blot analyses for CSN5 (D), CSN6 (E) and CSN8 (F). In-lane total protein images of the SDS-PAGE gel from the stain-free total protein imaging technology were used as the loading control. **G** and **H,** Representative images (G) and the summary of densitometry data (H) of western blot analyses for cytoplasmic and nuclear CSN5. **I** and **J**, Representative image (I) and pooled densitometry data (J) of native-PAGE followed by western blot for CSN5. Scatter plots superimposed by mean±SEM; each dot = a mouse (C-F) or a combined sample of 3 mice (A, B, G-J); unpaired Student’s *t*-test (B, H); two-way ANOVA followed by Tukey’s tests (D, E, F, J).

We next tested whether downregulation of CSN8 could affect other CSN subunits. CSN5 and CSN6 proteins in carotid arteries were also significantly reduced similarly to CSN8 in CSN8-hypo mice; the expression of CSN subunits were increased after vascular injury in the CTL mice, but this increase was not discernible in the CSN8-hypo mice (**Figure 4C-4F**).

Besides being an obligatory subunit of the CSN, each CSN subunit may participate in the formation of mini-complexes. CSN5 is unique in partitioning as a ∼500kDa holo-complex which is primarily located in the nucleus, and as mini-complex mostly localized in the cytoplasm.^15^ Nuclear fractionation revealed an increased protein level of CSN5 in the cytoplasmic fraction of the CSN8-hypo mice (**Figure 4G, 4H**). We further subjected the nuclear and cytoplasmic fractions to non-denatured gel electrophoresis followed by immunoprobing for CSN5, which revealed a significant increase in the abundance of CSN5 mini-complex located primarily in the cytoplasmic fraction and, concomitantly, a significantly decrease in the level of CSN5-contained holocomplex in both RCCA and the ligated LCCA of CSN8-hypo mice compared with control mice (**Figure 4I, 4J**). Taken together, these results indicate an essential role of CSN8 in stabilizing the holocomplex, as reduction of CSN8 alters the equilibrium of other CSN subunits in mouse arteries; and further demonstrate increased CSN5 mini-complexes in the cytoplasm of CSN8-hypo arteries, which is consistent with an augmented nuclear-export of p27, a likely mechanism underlying the enhanced VSMC proliferation and NH in CSN8-hypo mice upon vascular injury.

### 5. Increased proliferation and p27 nuclear exclusion correlating with elevated cytoplasmic CSN5 in cultured CSN8-hypomorphic VSMCs

Our *in vivo* findings have unveiled an exacerbation of NH by CSN8 hypomorphism in mice. We next tested if this could be VSMC-autonomous. VSMCs from the aorta of the CSN8-hypo and CTL mice were cultured and validated for the SMC identity using immunostaining and western blot analyses for SMC markers (**Figure S9**). To mimic a response to vascular injury in cell culture, we treated the cell with PDGF-BB, a potent mitogen that plays primary role in vascular response to injury and is known to induce VSMC proliferation and modulate phenotype switching.^32^ We found that PDGF-BB induced significantly more pronounced increases of PCNA in CSN8-hypo VSMCs compared with CTL VSMCs although, whereas no significant difference in VSMC proliferation was discerned between CSN8-hypo and CTL VSMCs under basal culture conditions (**Figure 5A, 5B**), recapitulating our in vivo findings.

**Figure 5.**
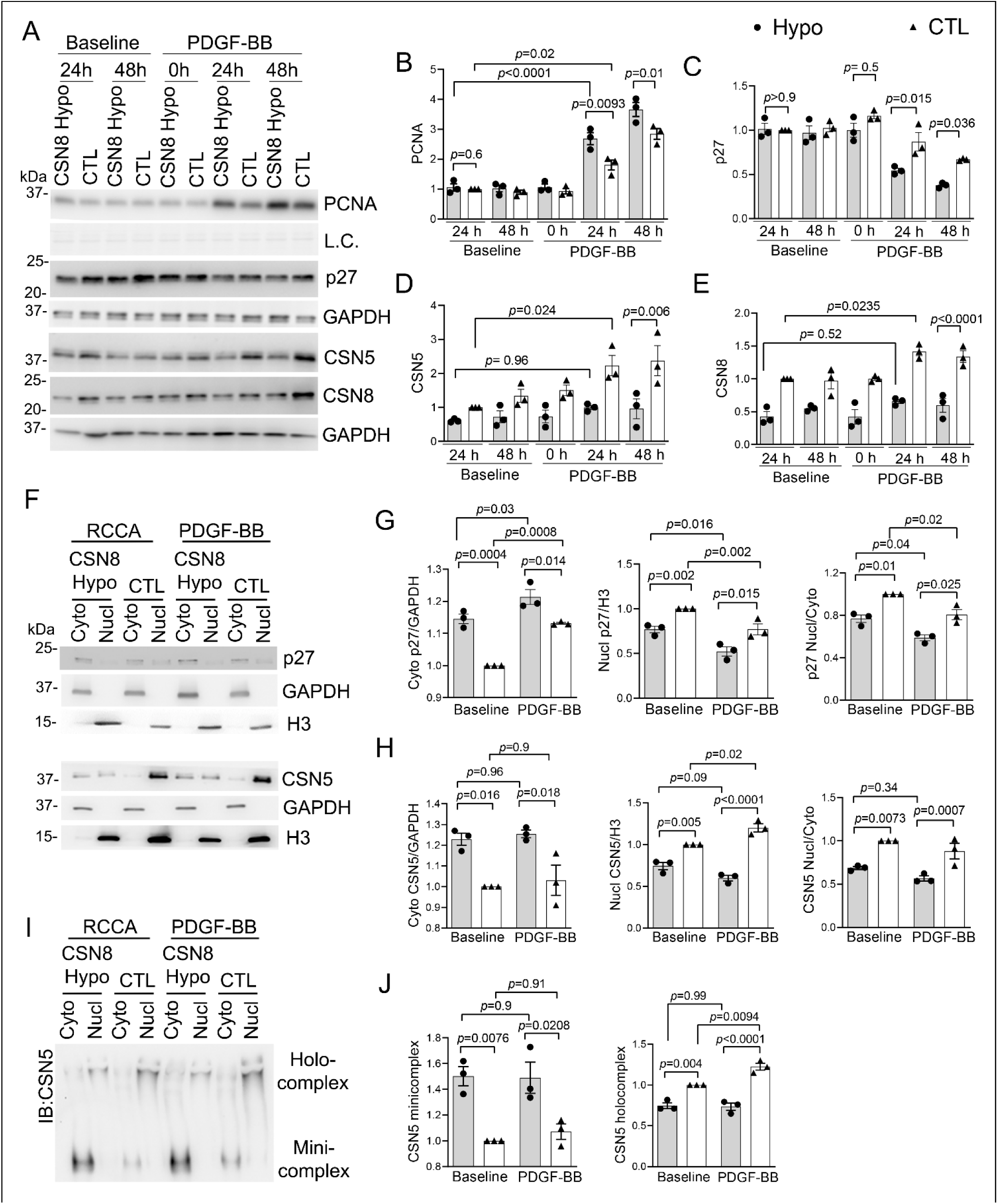
PDGF-induced cell proliferation and nuclear export of p27 and CSN5 are more pronounced in CSN8 hypomorphic VSMCs. Cultured VSMCs were serum-starved for 24 hrs, then treated with PDGF-BB or vehicle control for 24 or 48 hrs before collected for respective analyses. **A** ∼ **E**, Representative images (A) and pooled densitometry data (B through E) of immunoblot for PCNA (B), p27 (C), CSN5 (D), and CSN8 (E). **F** ∼ **H**, Representative image (F) and pooled densitometry data (G, H) of western blot analyses for cytosolic (Cyto) and nuclear (Nucl) p27 and CSN5. For western blot analyses, either GAPDH or stain-free total protein imaging was used as a loading control (L.C.). GAPDH and Histone H3 were probed as a marker and loading control for cytoplasmic and nuclear fractions, respectively. **I** and **J**, Representative image (I) and pooled densitometry data (I) of western blot analyses for cytosolic and nuclear native CSN5 protein complexes following native-gel electrophoresis. Mean±SEM; n=3 biological repeats; two-way ANOVA followed by Tukey’s tests.

In response to PDGF-BB stimulation, CSN8-hypo VSMCs displayed greater decreases in total p27 than the CTL VSMCs (**Figure 5A, 5C**). Additionally, subcellular fractionation revealed a strikingly decreased nuclear p27 and increased cytoplasmic p27 in the CSN8-hypo VSMCs compared to the CTL (**Figure 5F, 5G**). Consistent with our *in vivo* findings, protein levels of CSN5 and CSN8 were significantly decreased in hypomorphic VSMCs (**Figure 5A, 5D**, **5E**). Interestingly, the CSN subunits were upregulated upon PDGF-BB stimulation in control VSMCs (**Figure 5A, 5D**, **5E**). The effect of CSN8-hypo on CSN5 distribution was assessed using nuclear fractionation followed by native-PAGE and western blot for CSN5, which showed an increased prevalence of CSN5-minicomplex in the cytoplasmic fraction of CSN8-hypo VSMCs (**Figure 5I, 5J**). Together, these data demonstrate an increase of CSN5 minicomplex in the cytoplasm concurrently with a decrease of CSN holocomplex in the nuclei of CSN8-hypo VSMCs compared with CTL VSMCs, which correlates with increased nuclear export of p27 and subsequently increased VSMC proliferation.

### 6. Attenuation of the induced hyperproliferation of CSN8 hypomorphic VSMCs by pharmacological inhibition of nuclear export but not of CSN deneddylase

Our *in vivo* and *in cellulo* findings prompted us to further determine a mechanistic involvement for the nuclear export of p27 by CSN5 mini-complexes in the proliferation-promoting property of CSN8 hypomorphism. Leptomycin B (LMB) is a nuclear export inhibitor through blocking the interaction of CRM1 (chromosomal region maintenance 1, an export protein) with the nuclear export signal (NES) that functions to move proteins from the nucleus to cytoplasm (**Figure S10A**).^33^ CSN5i-3 is a CSN-specific deneddylase inhibitor,^18^ which is verified in cultured VSMCs (**Figure S10B**). Thus, we examined the effect of LMB (5 ng/ml) and CSN5i-3 (100 nM) on PCNA and p27 protein expression.

Treatment of VSMCs with LMB suppressed the PDGF-BB induced PCNA increases in both CSN8-hypo and control VSMCs (**Figure 6A, 6B**). Interestingly, CSN5i-3 significantly suppressed the PDGF-BB induced PCNA increases in control VSMCs but not in CSN8-hypo cells. LMB substantially diminished the difference in total p27 between the two genotypes (**Figure 6A, 6C**) and resulted in nuclear accumulation of p27 in both CSN8-hypo and control VSMCs (**Figure 6D, 6E**). CSN5i-3, however, did not alter the nuclear export of p27 (**Figure 6F**). These results indicate that blocking the nuclear to cytoplasmic shutting of p27 by inhibition of nuclear export abolishes the proliferation-promoting property of CSN8 hypomorphism in VSMCs and that the change in the nuclear export is independent of the deneddylase function of the CSN.

**Figure 6.**
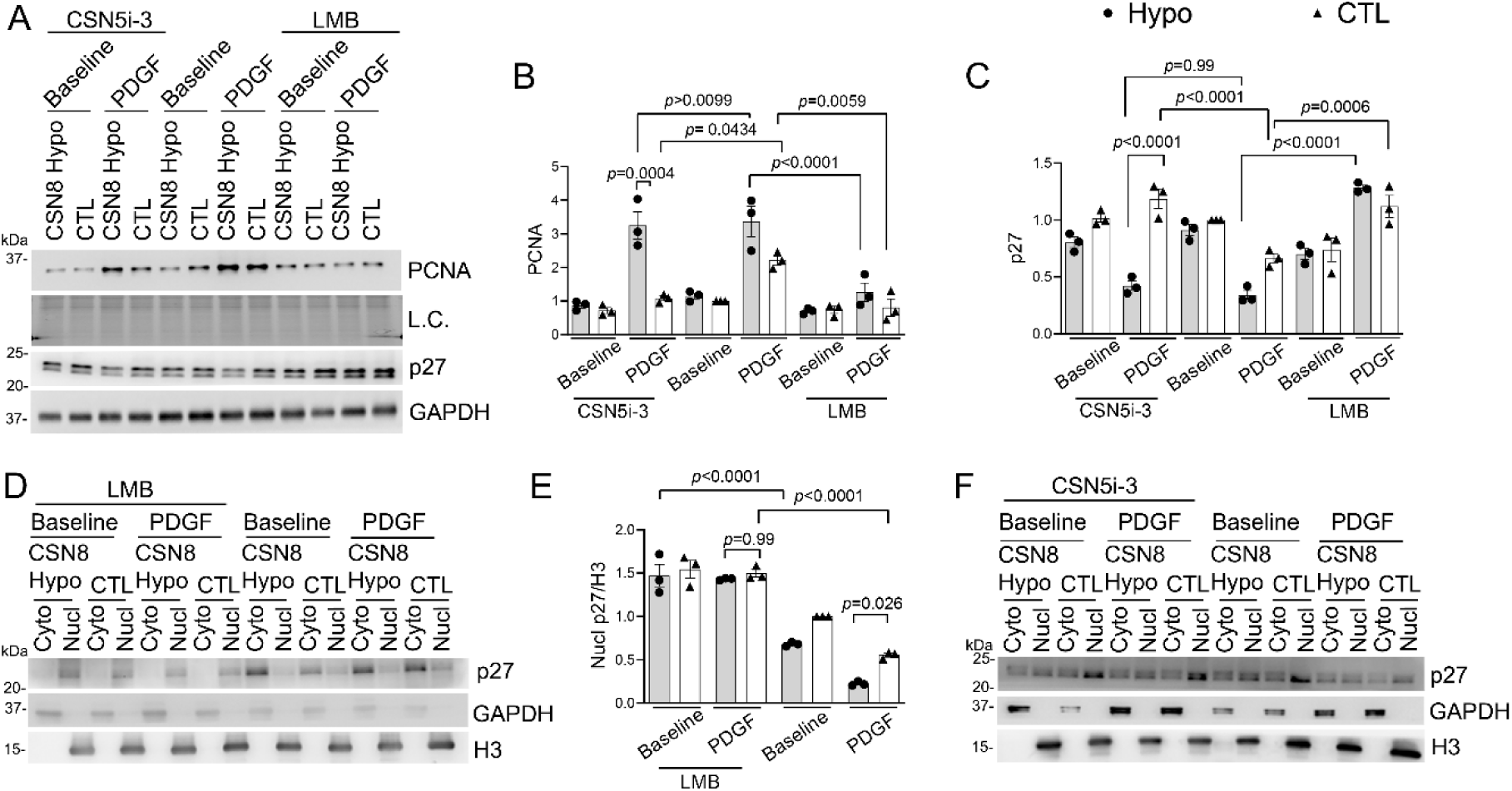
Nuclear export inhibitor leptomycin B (LMB) but not CSN deneddylase inhibitor CSN5i-3 suppresses the exacerbation of VSMC proliferation by CSN8 hypomorphism. VSMCs in cultures were treated with CSN5i-3 (100nM) or LMB (5ng/ml) in the absence or presence of PDGF-BB. Twenty-four hours after the treatment, cells were harvested for total protein extraction or nuclear fractionation. **A∼C**, Representative images (A) and pooled densitometry data (B, C) of western blot analyses for PCNA and p27. Loading control used either stain-free total protein imaging or GAPDH immunoblot images. **D** and **E**, Representative images (D) and the pooled densitometry data (E) of western blot analyses for nuclear (Nucl) and cytosolic (Cyto) p27 in VSMCs following LMB treatment. **F**, Representative images of western blot analyses for nuclear and cytosolic p27 in VSMCs treated as indicated. GAPDH and Histone H3 are used as loading controls and markers for cytosolic and nuclear fractions, respectively. Mean±SEM; n=3 biological repeats; two-way ANOVA followed by Tukey’s tests.

### 7. Restoring nuclear p27 and suppressing the hyperproliferation in CSN8 hypomorphic VSMCs by the nuclear export-disabled but not the deneddylase-dead CSN5

CSN5 contains the leucine rich NES which interacts with CRM1 and acts as an adaptor between p27 and CRM1 to mediate the cytoplasmic translocation of p27 in proliferating fibroblasts.^16^ To better understand the mechanistic role of CSN5 in VSMCs, we created full length CSN5 plasmids harboring mutation in either its NES or the MPN motif, referred to as the nuclear-export disabled CSN5 (CSN5-ΔNES) or the deneddylase-inactive CSN5 (CSN5-ΔMPN), respectively. Transfection of the mutant plasmids decreased the level of endogenous CSN5 in transfected VSMCs (**Figure 7A, 7F**), indicating that the transgenic CSN5 replaces the endogenous CSN5 in the CSN. Transfecting CSN5-ΔNES did not alter Cullin deneddylation (**Figure S11**) but significantly suppressed the PDGF-BB induced proliferation in both CSN8-hypo and control VSMCs; and abolished the difference in PDGF-BB-induced proliferation between the CSN8-hypo and control VSMCs as evidenced by the changes in PCNA protein levels (**Figure 7A, 7B**) and Ki-67 immunostaining (**Figure 7C**). A significant increase in the nuclear p27 was demonstrated following the transfection with CSN5-ΔNES (**Figure 7D, 7E**). Furthermore, PDGF-induced proliferation of control VSMCs was significantly suppressed by CSN5-ΔMPN, but the exacerbation of PDGF-induced proliferation by CSN8 hypomorphism (**Figure 7F, 7G**) and the subcellular localization of p27 (**Figure 7H**) remained unaffected. In addition, enforced overexpression of wild type (WT)-CSN5 significantly increased the proliferation of both CSN8-hypo and control VSMCs (**Figure S12**). Collectively, these experiments establish that CSN5-promoted nuclear export of p27 rather than CSN5-dependent deneddylation activity is responsible for the greater PDGF-BB induced proliferation in CSN8-hypo VSMCs.

**Figure 7.**
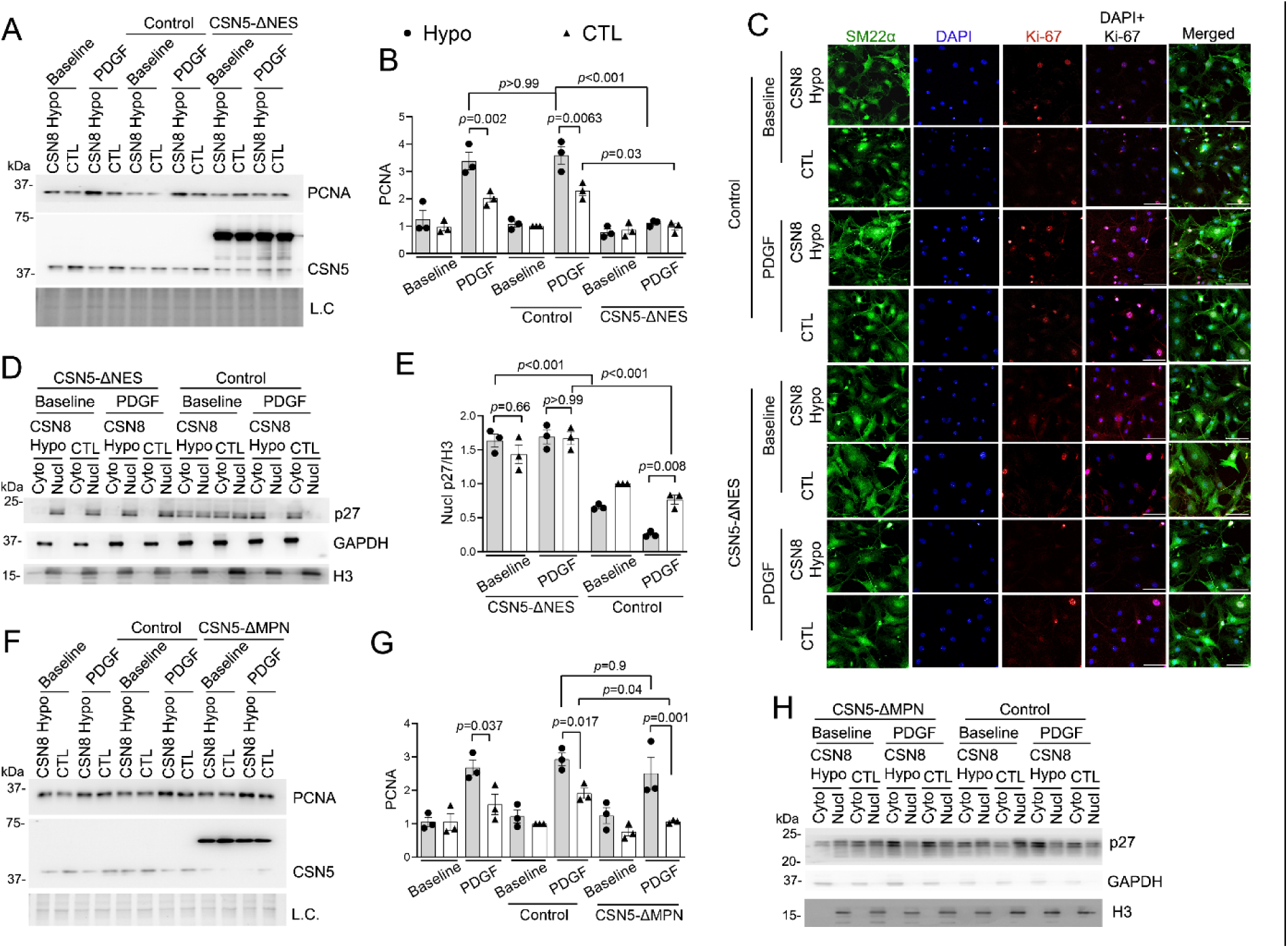
Overexpression of nuclear export-disabled CSN5 (CSN5-ΔNES) but not of deneddylase-dead CSN5 (CSN5-ΔMPN) restores nuclear localization of p27 and abolishes the hyperproliferation in CSN8 hypomorphic VSMCs. VSMCs were transfected with plasmids expressing EYFP-fused CSN5-ΔNES or CSN5-ΔMPN. The plasmids expressing EYFP alone were used as the control. The VSMCs were then treated with PDGF-BB or vehicle control for 24 h before harvested. **A** and **B**, Representative images (A) of western blot analyses for PCNA and CSN5 and pooled densitometry data of PCNA (B) in VSMCs treated as indicated. **C**, Representative confocal micrographs of immune-stained with Ki-67 (red) and SM22α (green) in VSMCs treated as indicated. Nuclei were stained with DAPI (blue). Scale bar= 100µm. **D** and **E**, Representative images (D) and the pooled densitometry data (E) of western blot analyses for nuclear (Nucl) and cytosolic (Cyto) p27 in VSMCs treated as indicated. F and G, Representative images (F) of western blot analyses for PCNA and CSN5 and the pooled densitometry data of PCNA (G) in VSMCs treated as indicated. **H**, Representative images of western blot analyses for nuclear and cytosolic p27 in VSMCs treated as indicated. For western blot analyses for PCNA, loading control (L.C.) used total protein stain-free imaging. GAPDH and Histone H3 are used as a loading control for cytosolic and nuclear protein, respectively. Mean±SEM; n=3 biological repeats; two-way ANOVA followed by Tukey’s tests.

### 8. Suppressing NH by pharmacological inhibition of CSN deneddylase in vivo

Intrigued by the findings with CSN5-SMKO and CSN5i-3 (*in cellulo*), we next examined the effect of CSN5i-3 *in vivo*. WT mice were treated with CSN5i-3 (20 mg/kg/day, i.p.) or vehicle control and subjected them to LCCA ligation. Ligated and control arteries were excised at the 7^th^ day of treatment and assessed for pharmacodynamics markers and the effect on VSMC proliferation and NH. The CSN5i-3 treatment regime did not show a significant effect on mouse body weight (*data not shown*). Neddylated Cul1 was increased, whereas Skp2 levels were reduced in the arteries isolated from the CSN5i-3 treated mice (**Figure 8A**). Administration of CSN5i-3 discernibly reduced the expression of PCNA in response to injury (**Figure 8B**). This was further corroborated by immunostaining for Ki67 and SM22α, which showed decreased VSMC proliferation and markedly attenuated NH in CSN5i-3 treated mice (**Figure 8C-8G**). Together, these data demonstrate that CSN5i-3 treatment effectively suppresses NH.

**Figure 8.**
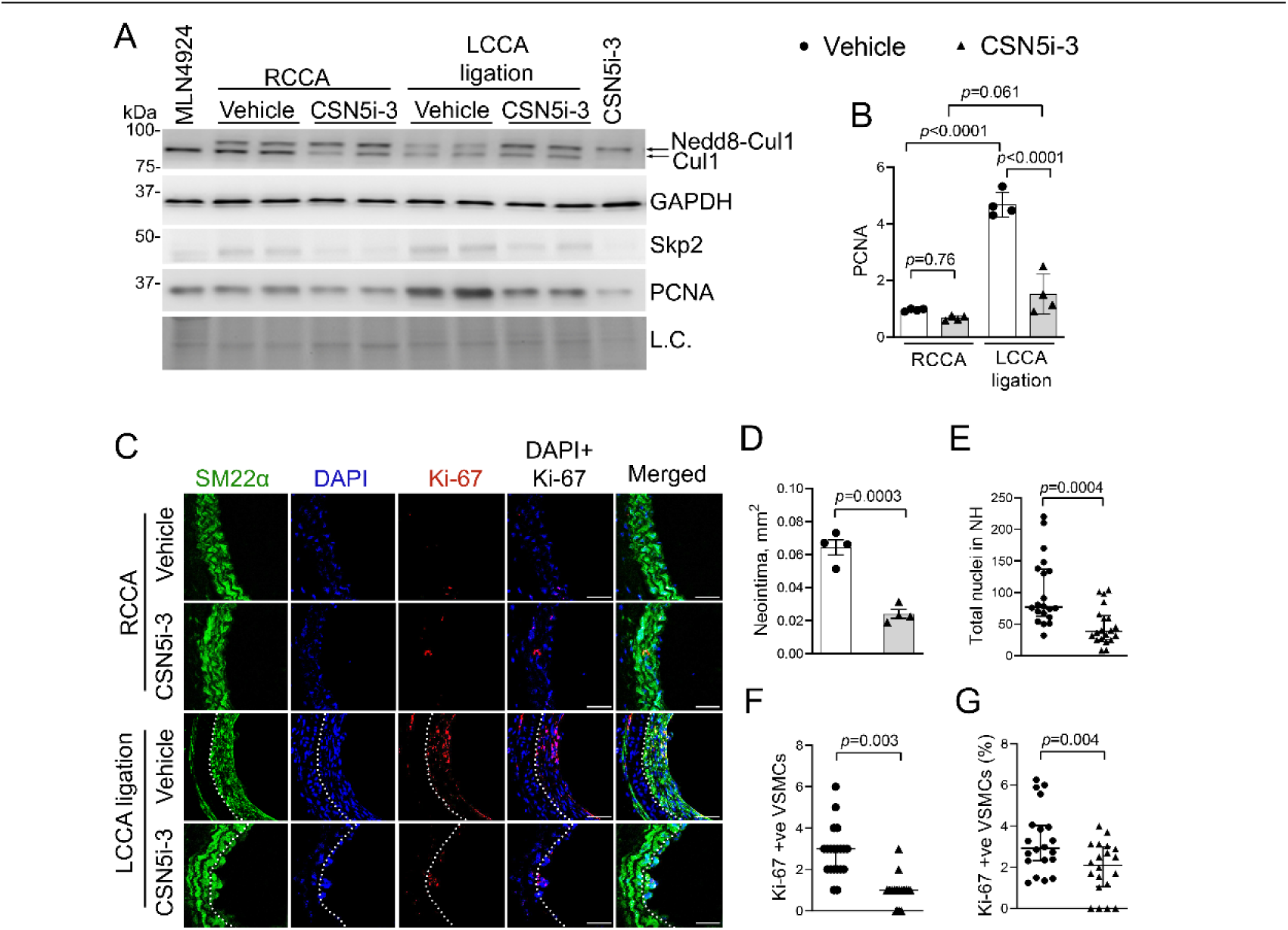
CSN5i-3 suppresses neointimal formation in vivo. Wild-type mice were administered with a dose of CSN5i-3 (20mg/kg, i.p.) or vehicle control 3 h before and daily after LCCA ligation. LCCA and RCCA were harvested 3 hours after the 7^th^ daily injection. **A** and **B**, Representative images of western blot analyses for indicated proteins (A) and pooled densitometry of PCNA (B). MLN4924 (neddylation inhibitor) and CSN5i-3 treated VSMCs were used to identify native and nedd8-Cul1 bands (the far-left and the far-right lanes in A), respectively. Scatter plots superimposed by mean±SEM where each dot represents a mouse and p values are derived from two-way ANOVA followed by Tukey’s tests. **C**, Representative fluorescence confocal images of immunostained Ki-67 (red) and SM22α (green) in the cryo-sections of LCCA and RCCA. Nuclei were stained with DAPI (blue). Scale bar= 50µm. **D,** Scatter plots superimposed by mean±SEM of the SM22α -positive neointima area in LCCA (2 mm proximal to the ligation site). Each dot represents a mouse; unpaired Students *t*-test. **E∼G**, Scatter plots superimposed by Median±95%CI of total number of nuclei in neointima (E), Ki-67 positive VSMCs in neointima (F), and the percentage of Ki-67 positive VSMCs (G) derived from microscopy as illustrated by panel C. Each dot represents a tissue section; 5 sections/mouse and 4 mice/group were included; Mann-Whitney test.

## DISCUSSION

Regulation of the cell cycle by the CSN and CSN subunits have been studied in several genetics systems,^34^ but the role of the CSN in VSMC function remains obscure. To our best knowledge, this is the first report demonstrating the *in vivo* significance of the CSN in VSMC proliferation and NH. Here we discover that vascular injury triggers the upregulation of the CSN and at least CSN5 in animals and humans, respectively. We further establish that both Cullin deneddylation by the CSN and the nuclear export exerted by CSN5 mini-complexes are responsible for the promotion of VSMC proliferation and NH by CSN5. Both CSN5-SMKO and pharmacological inhibition of the CSN deneddylase by CSN5i-3 effectively suppress VSMC proliferation and NH; and CSN8 hypomorphism exhibiting increased CSN5 mini-complexes exacerbates VSMC proliferation and NH. Thus, our study provides novel mechanistic insights into the CSN as a pivotal regulator of vascular remodeling, suggesting the CSN can be novel targets for prevention and treatment of NH.

### Upregulation of the CSN by vascular injury and the requirement of CSN deneddylase activity for neointimal hyperplasia

We discovered that CSN expression is upregulated in neointimal VSMCs in both animal models and human tissues. Both the mRNA and the protein levels of all examined CSN subunits as well as the CSN holocomplex were significantly increased in LCCA after ligation (**Figure 1**). Increased expression of CSN/CSN subunits were more often found in tumor tissues of multiple cancers.^34–36^ Complementarily to our findings, Asare *et al*. reported increases in CSN1, CSN5 and CSN8 expression in the endothelial layer of early human atherosclerotic plaques,^37^ whereas Liang *et al*. documented a critical role for loss of CSN6 at cardiac desmosomes in the pathogenesis of arrhythmogenic right ventricular dysplasia/cardiomyopathy.^38^ Our *in cellulo* data also reveal that the CSN subunits were upregulated in VSMCs stimulated with PDGF-BB (**Figure 5D, 5E**).

Comparable with our findings, a time-dependent upregulation of CSN6 was observed in cultured pulmonary arterial SMCs.^39^ Taken together, these findings suggest a pathogenic role for the CSN or its subunits in vascular remodeling. It will be interesting for future studies to determine the upstream events controlling CSN expression in the pathogenesis of vascular diseases.

The Cullin deneddylation activity of the CSN is critical to cell proliferation. In our study, we show that CSN5-SMKO accumulated neddylated Cul1 and Cul2 both at baseline and after LCCA ligation and attenuated the LCCA ligation induced increases in PCNA and Ki-67-positive VSMCs (**Figure 2**). Similar effects were achieved by pharmacological inhibition of the CSN by CSN5i-3 both *in cellulo* (**Figure S10B, 6A, 6B**) and *in vivo* (**Figures 8**). We reason that suppression of VSMC proliferation by impaired deneddylation acts through inactivating CRLs. It was purported that inactivation of CRLs by impaired Cullin deneddylation is because the sustained ubiquitination by the CRLs turns to destroy their own substrate receptor modules, resulting in accumulation of the substrate proteins of the CRLs.^18^ As a key substrate receptor for the CDK inhibitor p27 in the CRL1^SKP2^ ubiquitin ligase, SKP2 is essential for the ubiquitin-dependent degradation of p27 and thereby cell cycle progression.^17, 40, 41^ Here we detected that Skp2 proteins in vascular wall were markedly reduced by CSN5-SMKO (**Figure 2**) or CSN5-3i (**Figure 8**) and, reciprocally, total and nuclear p27 were significantly increased by CSN5-SMKO in LCCA wall (**Figure 2**) or by CSN5-3i in cultured PDGF-BB stimulated VSMCs (**Figure 6A**, **6C**), presenting a molecular link between loss of CSN deneddylase activity and the suppression of NH. Consistent with our findings, CSN5i-3 was reported to suppress cell proliferation and viability in tumor cell lines and inhibit growth of tumor xenografts in mice.^19, 42^

### Significant contributions of CSN5-mediated nuclear export to the promotion of neointimal hyperplasia by CSN5 and CSN8-hypo

Taking advantage of the increase of CSN5 mini-complex in CSN8-hypo cells where the deneddylation activity of the CSN is decreased and dissociated from other functions, the present study has unraveled a deneddylation-independent mechanism that governs SMC proliferation by CSN5. Prior study showed that CSN8 hypomorphic MEFs displayed an increased ratio of CSN5 mini-complex to the CSN holocomplex.^20^ Likewise, our study here reveals an increase in the abundance of cytoplasmic CSN5 mini-complex by CSN8 hypomorphism in VSMCs, accompanied by increased nuclear-export of p27 (**Figures 4, 5**). Importantly, NH and VSMC proliferation *in vivo* and *in cellulo* were exacerbated by CSN8 hypomorphism (**Figure 3, 5A**). Our data further demonstrate that enhanced nuclear-export by free CSN5 or CSN5 mini-complexes, independent of Cullin deneddylation, is responsible for the proliferation-promoting effect of CSN8 hypomorphism. First, despite a reduction in protein levels of CSN5 and CSN6 by CSN8 hypomorphism (**Figure 4C-4F**), which is consistent with the notion that ablation of any individual CSN subunits disrupts the CSN holocomplex and results in loss of the Cullin deneddylation activity and destabilization of remaining CSN subunits,^20, 31^ cytoplasmic CSN5 mini-complexes were significantly increased by CSN8-hypomorphism (**Figure 4I**, **4J**, **5I, 5J**). Second, neither pharmacological nor genetic inhibition of CSN deneddylase activity could rescue the reduced nuclear p27 and the increased proliferative phenotype in CSN8 hypomorphic VSMCs (**Figures 6, 7**). And lastly, both inhibition of nuclear export by LMB and genetically disabling the nuclear-export function of CSN5 restored nuclear p27 levels and attenuated the CSN8 hypomorphism-induced VSMC hyperproliferation (**Figure 6, 7**). Corroborating our findings, Tomoda *et al*. reported that CSN5 served as an adaptor for shuttling p27 from the nucleus to the cytoplasm of proliferating fibroblasts.^16^ Notably, although the CSN subunits were upregulated after vascular injury and PDGF-BB stimulation in controls, CSN8 hypomorphism prevented this upregulation (**Figure 4C-4F**, **5A, 5D, 5E**) but exacerbated NH and VSMC proliferation (**Figures 3, 5**). Previously, CSN8 was shown to be necessary for the cell cycle re-entry in periphery T cells and MEFs and essential for postnatal hepatocyte survival and effective proliferation.^20, 43, 44^ Surprisingly, though CSN8 hypomorphism showed increased p27 and CSN5 in the cytoplasm, there was no difference observed in terms of proliferation in baseline, suggesting that an increase in CSN5 mini-complex is not enough to initiate cell proliferation but is capable of promoting the proliferation process by, for example, shortening the G1 phase.^20^

### Targeting CSN5 to treat neointima hyperplasia

NH is prominent in the pathology of primary PAH, a rare but devastating disease that lacks effective clinical treatment beyond lung transplantation.^8^ Restenosis due to NH remains a prevailing clinical problem and aberrant VSMC proliferation threatens almost every known vascular reconstructive procedure, bringing new cardiovascular risk to postoperative patients.^4, 6^ The mechanism underlying this proliferative process is not completely elucidated, hindering the search for effective countering measures. The present study demonstrates not only that genetic ablation of CSN5 in SMCs inhibits VSMC proliferation and NH in animals (**Figure 2**) but also that pharmacological inhibition of the CSN by CSN5i-3 exerts striking suppression on VSMC proliferation *in cellulo* and in vivo (**Figure 6, 8**) and on NH in animals (**Figure 8**), providing extremely promising support for targeting the CSN to treat NH-based disorders, such as PAH and restenosis. For collecting proof-of-principle evidence, present study only tested one effective regime of CSN5i-3 to suppress VSMC proliferation and NH after LCCA ligation, where the treatment is well tolerated although moderate cardiotoxicity was observed at the end of treatment regime (*data not shown*). This is in agreement with the more severe phenotype caused by cardiomyocyte-restricted knockout of CSN8 in mice.^31^ Hence, it will be important to further test the effectiveness of lower degrees and shorter durations of pharmacological inhibition of the CSN for clinical translation. To this end, it will be extremely interesting to test the effect of local and controlled administration of the inhibitor(s) via, for example drug-eluting stents on post-operative restenosis.

## Acknowledgements

We thank Megan T. Lewno, Jose R. Lira, and Jack O. Sternburg for their outstanding technical assistance in managing mouse colonies and genotyping for this study.

## Supplemental material

Supplemental methods

Supplemental Table I and II

Supplemental Figures S1-S12

Supplemental References ^45–47^

## Non-standard Abbreviations and Acronyms

CDK: cyclin-dependent kinase
CRLs: Cullin-RING ligases
CSN: the COP9 signalosome
CTL: the control group
Cul: cullin
KO: knockout
LMB: leptomycin
NES: nuclear export signal
NH: neointimal hyperplasia
PDGF: platelet-derived growth factor
PCNA: proliferating cell nuclear antigen
UPS: ubiquitin-proteasome system
VSMC: vascular smooth muscle cell
WT: wildtype

