## Supplemental methods and data for "COP9 Signalosome Promotes Neointimal Hyperplasia via Deneddylation and CSN5-Mediated Nuclear Export"

### I. A Full Description of Materials and Methods

#### Animal models

CSN5 (also known as JAB1) is encoded by Cops5. Mouse with a Cops5-floxed allele contains exon 2 flanked by 2 LoxP sites inserted in the introns sandwiching exon 2. Cre-mediated recombination effectively removes the exon 2, resulting in an early frameshift and translation termination.[25] The Myh11-CreERT2 transgenic mouse was obtained from Jackson Laboratory, # 019079 (B6.FVB-Tg (Myh11-icre/ERT2)1Soff/J).[26] The generation of CSN5$^{flox/flox}$::Myh11-CreERT2 was achieved though crossbreeding Myh11-CreERT2 transgenic mice with CSN5$^{flox/flox}$ in C57BL/6 genetic background. Further, to achieve homozygous smooth muscle-restricted CSN5 knockout (CSN5-SMKO) in adult mice, we treated CSN5$^{flox/flox}$::Myh11-CreERT2 mice with tamoxifen, with the Myh11-CreERT2 mice as a control. The mice were routinely checked after the inducible knockout. The Myh11-CreERT2 transgenic mouse harbors in its Y-chromosome a transgenic cassette that expressed a mutant estrogen receptor sandwiched Cre recombinase (CreERT2) under the control of the promoter of the myosin heavy chain 11 (Myh11); the latter is SMC-specific. Hence, limited by the Myh11-CreERT2 transgene insertion location (Y chromosome), only male mice were suitable for this part. So far, no reported studies have investigated CSN5$^{flox/flox}$::Myh11-CreERT2.

Generation of CSN8/Cops8 targeted alleles has been previously described.[20] Briefly, Cops8$^{neoflox}$ allele contains a neomycin resistant cassette in intron between exon 3 and 4 while CSN8 knockout allele (CSN8$^{-}$ ) has a deletion of exon 4 to 6. The C57BL/6J Cops8$^{neoflox/+}$ and Cops8$^{+/-}$ mice were backcrossed into FVB/N background for at least six generations. The homozygous Cops8$^{neoflox/neoflox}$ mice were then mated with Cops8$^{+/-}$ mice to produce Cops8$^{neoflox/-}$ and Cops8$^{neoflox/+}$mice, which were used as CSN8 hypomorphic mice and control mice, respectively. The neomycin gene inserted in an intron of a CSN8-floxed allele (CSN8$^{neoflox/+}$) reduces CSN8 gene expression but this reduction of CSN8 gene expression does not cause CSN8 protein reduction until the other CSN8 allele is deleted (CSN8$^{neoflox/-}$); hence, the CSN8$^{neoflox/-}$ mice have been confirmed as CSN8 hypomorphic (CSN8hypo) and the littermate CSN8$^{neoflox/+}$ mice were used as controls (CTL).

All procedures involving animals were approved by the Animal Care and Use Committee of the University of South Dakota and conform to the NIH Guide for the Care and Use of Laboratory Animals.

#### Tamoxifen administration for smooth muscle cell restricted CSN5 knockout mice (CSN5-SMKO).

To achieve homozygous CSN5-SMKO in adult mice, we treated CSN5$^{flox/flox}$::Myh11-CreERT2 mice with tamoxifen (Catalog#HY-70062; MedChem Express, Monmouth, NJ) with the Myh11-CreERT2 mice as a control. Tamoxifen was dissolved in 100% ethanol through extensive vortexing and mild heating (< 40°C) to solubilize completely. The tamoxifen-ethanol solution was mixed with autoclaved sunflower oil at the final concentration of 10ug/ul and stored in light blocking vessel. The tamoxifen-oil mix (1mg tamoxifen/mouse) was intraperitoneally (IP) injected in adult mouse. Each animal went through 2 rounds of IP for 5 days with 2 days break between, with total of 10 injections.

Mice were closely monitored for any adverse reaction to the treatment. After the last IP, we waited for at least 2 weeks to wash off the side-effects of exogenous tamoxifen.

**Ligation of the left common carotid artery (LCCA)**

To produce NH in vivo, LCCA ligation was performed on adult mice as described.[27] Briefly, mice were anesthetized using 2.5% isoflurane, the left carotid bifurcation was located and then ligated with 6.0 silk suture immediately approximal to the bifurcation. The mice were monitored until recovery and buprenorphine SR was used as an analgesic. Animals were then observed each day until tissue collection for pain or any health-related problems. After certain time after ligation, the mice were sacrificed and the ligated LCCA section was collected for further processing. The right common carotid artery (RCCA) was used as the uninjured intra-animal control vessel.

**Cell culture**

VSMCs were enzymatically isolated from abdominal aortas of adult mice as described.[28] Briefly, mice were euthanized using a continuous flow of carbon dioxide and the skin was opened to expose the abdomen and thorax using surgical scissors. The entire abdominal aorta up the renal bifurcation was dissected and cleaned off the surrounding fat and adventitial tissue. The cut pieces of aorta were digested in collagenase II for 6 hours at 37°C and then were grown in DMEM supplemented with 10% fetal bovine serum and 1% antibiotic/antimycotic solution at 37°C in a humified atmosphere of 5% CO2. VSMCs at passage 3-6 were used for all experiments. Before each experiment, the cells were serum starved and then treated with PDGF-BB, 10ng/ml (Catalog#220-BB; R&D biosystems, Minneapolis, MN) or the vehicle control (PBS).

**Immunohistochemical assessment of CSN5 in human pulmonary arteries**

Human lung tissue samples were taken from archived surgical pathology paraffin embedded tissue acquired as standard of care. The control tissue represents sections of normal segment of lung tissue taken during surgical resection of lung tumors. The PAH tissue was acquired from explanted lungs from patients with idiopathic PAH undergoing transplantation. The demographic information of the patients is shown in **Supplemental Table II**. Immunohistochemistry detection was performed with VitroView Universal 1-step polymer-based IHC/DAB kit (Catalog# VB-6023D, VitroVivo Biotech, Rockville, MD) as per manufacturers instruction with minor modification. Paraffin embedded sections were deparaffinized with xylene (3X5min) and rehydrated through series of ethanol (100% ethanol for 2X2mins, 95% ethanol for 2x2mins, 70% ethanol for 2mins and 50% ethanol for 2 mins) prior to histological staining. Antigen retrieval was performed by microwaving the sections in citrate buffer (10mM, pH 6) with 0.05% Tween-20 for 15 mins. Paraffin sections were then permeabilized using 0.3% Triton X-100 for 10 mins, washed with 1XPBS, and blocked for non-specific binding of immunoglobin in normal goat serum for 1h. Afterwards, the sections were incubated in mouse anti-CSN5 (Catalog#sc-13157, Santa Cruz, Dallas, Texas) overnight at 4°C. For chromogenic detection, sections were blocked in 0.3% hydrogen peroxide for 10 mins at room temperature; and the primary antibody detection was performed using polymeric peroxidase anti-mouse/rabbit secondary antibody for 1h at room temperature, followed by colorimetric detection using DAB substrate. The slides were then washed using distilled water and counterstained with hematoxylin solution as per manufacturers instruction. The sections were dehydrated though series of ethanol (70% ethanol for 2 mins, 95% ethanol for 2 min and 100% ethanol for 2X3min), cleared in xylene (2X5min) and finally mounted using Permount (Fisher chemical). Negative controls were performed with the omission of the primary antibody. Images were viewed by light microscope (Zeiss AxioImager M1, Taunton, MA) and for quantitative readouts, Fiji Image J (plugin: color deconvolution, IHC image analysis toolbox) was used using methods as previously described.[45] In brief, the image was segregated to obtain individual images for hematoxylin and DAB staining. The threshold was adjusted to convert the brown image into a black and white binary mask image. The area for measurement (tunica media for control and the neointima for PAH sections) were lined and the stained positive area was calculated. Similarly, the segregated hematoxylin image was adjusted for threshold, converted to binary image, and overlaid with the positive DAB staining to calculate the CSN5 positive nuclei. For each sample slide, we assessed 3 different pulmonary arterial sections representing different fields.

**Protein extraction and western blot analysis**

The surrounding adventitia of the collected carotid arteries was gently removed. Collected arteries or cultured VSMCs were lysed in lysis buffer (41 mM tris-HCl, 1.2% SDS, and 8% glycerol), subsequently sonicated, boiled for 5 mins and centrifuged at 10,621 x g for 10 mins at 4°C. The supernatant was collected for western blot analyses and protein concentration was quantified using the bicinchoninic acid assay (BCA). Equal amounts of protein were loaded in SDS-polyacrylamide gel (8-16%), transferred to polyvinylidene difluoride (PVDF) membrane, and incubated with primary antibodies against the protein of interest. The incubated PVDF membrane was washed to remove unbound primary antibodies, followed by the incubation with horseradish peroxidase (HRP) conjugated secondary antibodies and again, washed to remove unbound antibodies. The bound secondary antibodies were detected using the enhanced chemiluminescence detection reagents. Blots were imaged and quantified using the Image Lab software (Bio-Rad). Either GAPDH or the total protein content derived from the stain-free protein imaging technology was used as in-lane loading control. The antibodies used are detailed in **Supplemental Table I**.

**Cytoplasmic and Nuclear fractionation**

Cytoplasmic and nuclear fractions were extracted using the Epiquik nuclear extraction kit (Catalog #OP-0002-1; Epigentek, Farmingdale, NY) according to the manual provided by the manufacturer. For tissues, 3 carotid arteries were combined per group. In brief, tissues or cultured cells were washed or collected using PBS and centrifuged to collect the pellet. The pellet was then suspended in cytoplasmic extraction buffer, incubated in ice for 10 minutes after which it was vortexed vigorously for 10 sec and finally, centrifuged to collect the supernatant, cytoplasmic fraction. The remaining pellet was then washed with PBS two times, suspended in nuclear extraction buffer, and incubated on ice for 15 mins (vortexing the tube every 3 mins). The tube was then centrifuged for 10 mins at 20,817 x g and the supernatant, nuclear fraction was collected. Both cytoplasmic and nuclear fractions were stored at -80°C until further use.

GAPDH and Histone H3 were used as a marker for cytoplasmic and nuclear fractions respectively.

**Native Gel Electrophoresis**

Total protein isolated from carotid arteries, or the homogenates isolated using nuclear and cytoplasmic kit were used for running native gel. For the former, 2 carotid arteries were combined per group. The tissue was then homogenized in an extraction buffer (50mM Tris-HCL, pH 7.5, 1mM ATP, 5mM MgCl2, 1mM DTT, 250 mM Sucrose) and centrifuged at 4°C for 30 min (15000 x g). Protein concentration was determined with BCA reagents and the samples were diluted with 4X native gel loading buffer [200mM Tris-HCL, pH 6.8, 60%(v/v) glycerol, 0.05%(w/v) bromophenol blue]. Equal amounts of protein (20ug) were separated on a 4% gradient gels at 100V and 4°C in the SDS-free running buffer for about 3-4 hours. Following, conventional western blot analysis was performed using antibodies specific for indicated CSN subunits.

**RNA isolation, cDNA synthesis and quantitative PCR**

Total RNA was extracted from ligated LCCA and unligated RCCA from wildtype mice using the TRI Reagent (Molecular Research Center Inc., Cincinnati, OH). The concentration of RNA was determined using Agilent RNA 6000 Nano assay (Agilent technologies Inc., Germany) following the manufacturer's instruction. For reverse transcription (RT) reaction, 1 µg of RNA was used as a template to generate complementary DNA using the high-capacity cDNA reverse transcription kit (Catalog #4368814; ThermoFisher Scientific, Waltham, MA), and the RT was performed by following the manufacturer's instructions. Transcripts of interest were visualized and quantified using conventional semi-quantitative PCR (RT-PCR) and quantitative real-time PCR (qPCR) with the SYBR-Green assay respectively. For the conventional RT-PCR, 2 µl of solution from the RT reaction and specific primers toward the target gene and GAPDH were used. The mRNA levels of the gene of interest were visualized by PCR at the minimum number of cycles (15 cycles) capable of detecting the PCR products within the linear amplification range.

Further, the relative quantification of mRNA expression was carried out using ~100ng of cDNA per reaction. qPCR was performed in technical duplicates in 20ul of reaction volume containing 200nM of specific primers and 5 ul of SYBR using real-time detection system. Level of gene expression was normalized to internal control, GAPDH, calculated as $\Delta$Ct and $\Delta\Delta$Ct, and plotted as $2^{-\Delta\Delta Ct}$ (fold-change). Mouse CSN8, CSN5, CSN6 and GAPDH primer sequences are as follows: CSN8, 5`-GTCAGTTGGACAGCGAATCT-3` (forward) and 5`-CGTCTCCTTHTTGCATCTCTAA-3`(reverse); CSN5, 5`-CCCTCCTTCTACCTTGTTTGAG-3` (forward) and 5`-CTGTCGATGGTCAGATGAGAAG-3`(reverse); CSN6, 5`-TTCCTGGGCTTTGCCTTATC-3` (forward) and 5`-GGGTCAACAACACTCACATCT-3` (reverse); GAPDH, 5′-ATGACATCAAGAAGGTGGTG-3′ (forward) and 5′-CATACCAGGAAATGAGCTTG-3′ (reverse).

**Histological processing and immunostaining of mouse carotid artery sections**

Carotid artery sections for Hematoxylin and Eosin (H&E) staining were embedded in paraffin blocks. The tissue was cut to obtain 5μm sections, placed on glass slides, air dried and heat fixed overnight. Sections were deparaffinized and rehydrated through xylene and series of decreasing alcohols. Following, the sections were stained with hematoxylin and rinsed with tap water. After counterstaining with eosin, the sections were dehydrated and mounted with xylene. The imaged sections were then visualized (Olympus IX71) and analyzed using Image J for morphometric parameters. Immunohistochemical assessment of CSN5 in mouse tissue was processed similarly as of human paraffin lung sections as mentioned above.

Immunofluorescence staining and confocal microscopy were performed on cryosections from ligated LCCA, unligated RCCA or cultured VSMCs. 4% paraformaldehyde-fixed VSMCs on cover glass or tissue cryosections (7μm) were washed thrice for 5 min with PBS, incubated with 1% glycine in PBS for at least 30 min. The tissue or cells were incubated with 0.1% Triton-X-100 for 10-15 mins and subsequently, blocked with 2% BSA at room temperature for 1 h. Primary antibodies were then added to specimen and incubated overnight at 4°C. Unbound antibodies were removed via 5 min x 3 washes with PBS before incubation with appropriate secondary antibodies at room temperature for 1 h. The specimens were then rinsed with PBS 3 times for 5 min each. DAPI was used for staining nuclei. The stained sections were covered by glass coverslips, sealed with nail polish, and kept in -20℃ prior to confocal imaging analyses. The fluorescence staining was visualized and imaged using a confocal microscope (Leica DMi8). Ligated and unligated carotid arteries were stained for CSN5 or CSN8 and SM22α to examine the expression of the CSN in neointima VSMCs. Further, carotid artery and cultured VSMCs were analyzed for SM22α, DAPI and Ki-67 positive VSMCs; for ligated LCCA, 5 sections per mouse were stained and the colocalization between the SM22α and Ki-67 with the nuclei (DAPI) in the intimal area was examined. Immunofluorescence imaging was collected and processed similarly in the experimental and control groups. To minimize non-specific binding, the primary antibodies (**Supplemental Table I**) was used at an optimized concentration for the immunostaining.

**Plasmids**

EYFP-JAB1 was a generous gift from Johannes A. Schmid (Addgene plasmid # 111213; http://n2t.net/addgene:111213; RRID: Addgene_111213).[46] Plasmid amplification was carried out using maxiprep/midiprep/miniprep kit in accordance with the manufacturer's instruction. Restriction enzymes, BamHI (Catalog# R0136S; New England Biolabs, Ipswich, MA) and HINDIII Catalog# R0104S; New England Biolabs, Ipswich, MA); and T4 DNA ligase (Catalog# M0210S; New England Biolabs, Ipswich, MA) were used in accordance with the manufacturer's protocol. Transformation into *E. coli* was achieved with Subcloning Efficiency DH5α chemically competent cells (Catalog# 18265017; ThermoFisher Scientific, Waltham, MA) according to manufacturer's instruction.

The mutants were constructed by a three-step site specific PCR approach. To achieve the constitutive-nuclear CSN5, primers containing mutation sites against leucine residues at 237,238, and 240 were strategized and converted to alanine.[16] The primers were used at a final concentration of 200nM. In the first step, a mutagenic primer was used as the 3` primer and external primer was used as the 5` primers. The latter 5` external primer contains the BamHI recognition site for cloning. This step produced a mutant product. In the second step, a second PCR was carried out with the mutagenic primer as the 5`primer (complement of the first mutagenic region) and an external primer at the 3` regions. The 3`end contained the HINDIII recognition site for cloning. After each PCR step, the mutant product was run on an agarose gel to confirm the presence of desired base pair of PCR product. A fusion overlapping PCR was then carried out to combine the above two PCR products (200ng of each) with the end primers BAMHI and HINDIII in order to obtain the desired CSN5 mutant. The primers were added after nine cycles of fusion PCR in order to prevent the amplification of multiple PCR products due to traces of primers present in the template. The CSN5 mutant product was then TA cloned (Catalog# K202020; ThermoFisher Scientific, Waltham, MA) following the manufacturers instruction and sequenced (sangers sequencing) to confirm the presence of desired mutation. The mutant construct (insert) was digested using restriction enzymes, and gel purified. The mutant CSN5 insert was then ligated into EYFP vector containing the CMV promoter using the T4 DNA ligase, transformed, and amplified. The orientation and ligation were reconfirmed by restriction digestion using BAMHI and HINDIII as recognition sites.

Similarly, for the deneddylase-dead CSN5, histidine residues at 140 and 142 from the protypical MPN-JAMM consensus sequence $EX_n HSHX_7SXXD$[47] were targeted for mutation and converted to aspartic acid using strategy as mentioned above. The mutants were constructed using the primer pairs listed below.

CSN5-ΔNES

Forward 5`-CACCATCTTCCGCCGAGCCGATGCTGA-3`

Reverse 5`-TCAGCATCGGCTCGGCGGAAGATGGTG-3`

CSN5-ΔMPN

Forward 5`- GCCAGGGTCGCTATCATACCACCCGA-3`

Reverse 5`-TCGGGTGGTATGATAGCGACCCTGGC-3`

External primers

BAMHI 5`- GATCCGGTGGATCCGTTAAGAG-3`

HINDIII 5`-TCGAGCTCAAGCTTCCATGGC-3`

The control EYFP was prepared by gel cutting the backbone obtained by restriction digestion with BAMHI and HINDIII. Following the 3` and 5` end were blunted using the Klenow fragments (Catalog# M0210S; New England Biolabs, Ipswich, MA) as per manufacturers instruction. The ends were ligated and further amplified.

**Cell transfection**

Transient transfection was achieved with lipofectamine 3000 transfection reagent (Catalog# L3000001; ThermoFisher Scientific, Waltham, MA) using manufacturers protocol. In brief, cells were plated until it reached 70-80% confluency. Lipofectamine

3000 was diluted in Opti-MEM medium. Then, a master mix was prepared by diluting DNA in Opti-MEM medium and the enhancer reagent, P3000TM was added, mixed, and incubated at room temperature for 15-20 mins. The DNA-lipid complex was then added to the cells and checked for the transfected cells 48-72 post-transfection.

**CSN5i-3 administration *in vivo***

CSN5i-3 (Catalog# HY-112134; MedChem Express, Monmouth, NJ) was formulated in 90% autoclaved corn oil and 10% DMSO at a concentration of 20 mg/kg. Starting, young mice were injected with CSN5i-3 intraperitoneally and 3 h of the first injection, ligation of LCCA was performed. Following, mice were injected each day until the last day of treatment and closely monitored for any health-related issues or overt phenotype. At day 7, post 3 h of the injection, samples were collected for respective experiments.

**Criteria for selection of representative image**

The selection of the representative image was based on: (1) the image was free of technical defects and obtained with an optimal exposure time, (2) the image best represents the central tendency of the group.

**Statistical analysis**

Data were statistically analyzed using GraphPad Prism (version 9, GraphPad Software, San Diego, CA). All quantitative data are presented as mean±SEM unless otherwise indicated. Differences between the two groups were evaluated using two-tailed unpaired Student's *t* test and, when difference between three or more groups were evaluated, one-way or, where appropriate, two-way analysis of variance (ANOVA) followed by Tukey's multiple comparison test was used. For non-parametric data sets, Mann-Whitney test was also used. Statistical tests used to assess the statistical significance are indicated in each figure legend and the p-value is provided in the graphs. The *p* value or adjusted *p*-value $<0.05$ was considered statistically significant.

**Supplemental Table I. Primary/secondary antibodies used in this study.**

| **Target antigen** | **Vendor/Source** | **CATALOG #** | **Working concentration** |
|---|---|---|---|
| PCNA | Abcam | Ab15497 | 1:1000 for WB |
| P27 | Cell signaling | 3686 | 1:1000 for WB |
| CSN8 | Enzo | BML-PW8290-0100 | 1:1000 for WB<br>1:200 for IF |
| CSN5 | Bethyl | A300-014A | 1:1000 for WB<br>1:100 for IF |
| CSN6 | Enzo | BML-PW8295 | 1:2000 for WB |
| CSN4 | Bethyl | A300-013A | 1:1000 for WB |
| CSN3 | Bethyl | A300-012A | 1:1000 for WB |
| CSN1 | Bethyl | A200-026A | 1:1000 for WB |
| GAPDH | Santa Cruz | Sc-32223 | 1:1000 for WB |
| Histone H3 | Abcam | Ab18521 | 1:1000 for WB |
| Cullin1 | Santa Cruz | Sc-17775 | 1:1000 for WB |
| Cullin2 | ThermoFisher | 51-1800 | 1:2000 for WB |
| Desmin | Abcam | Ab15200 | 1:1000 for WB |
| TAGLN/Transgelin/ SM22α | Abcam | Ab14106 | 1:1000 for WB<br>1:400 for IF |
| Ki-67 | ThermoFisher | 14-5698-82 | 1:500 for IF |
| SKP2 | Cell signaling | 4358 | 1:500 for WB |
| Anti-YFP | Abnova | MAB8759 | 1:1000 for WB |
| 488-Anti Rabbit | Jackson Immunoresearch | 111-545-003 | 1:500 for IF |
| 647-Anti-Rat | Jackson Immunoresearch | 112-545-003 | 1:500 for IF |
| 647-Anti-Rabbit | Jackson Immunoresearch | 111-605-003 | 1:500 for IF |
| 647-Anti Mouse | Jackson Immunoresearch | 115-605-003 | 1:500 for IF |
| Anti-Rabbit HRP | Jackson Immunoresearch | 111-035-003 | 1:1000 for WB |

| | | | |
|---|---|---|---|
| Anti-Mouse HRP | Jackson Immunoresearch | 111-035-003 | 1:1000 for WB |
| Anti-Goat HRP | Jackson Immunoresearch | 305-035-003 | 1:1000 for WB |

#### II. Supplemental Data

**Supplemental Table II. Demographic information of the patients for the human lung tissue samples with pulmonary arterial hypertension (PAH) and without PAH (CON).**

| PATIENTS | AGE | GENDER |
|---|---|---|
| CON01 | 73 | F |
| CON02 | 68 | F |
| CON03 | 71 | F |
| PAH01 | 52 | F |
| PAH02 | 50 | F |
| PAH03 | 25 | F |

F, Female.

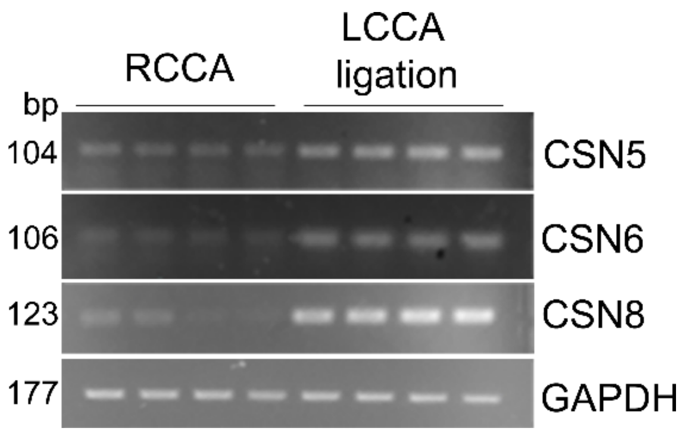


**Figure S1. The mRNA levels of CSN subunits are upregulated in wildtype mice after ligation of the left common carotid artery (LCCA).** Adult wildtype mice were subject to LCCA ligation; 1 week later, the LCCA segment proximal to the ligation site as well as the corresponding segment of the right common carotid artery (RCCA) were collected for RNA extraction. Shown are representative image of conventional reverse transcriptase PCR (RT-PCR) of indicated CSN subunits, the result is consistent with the real time PCR (qPCR) results shown in Figure 1C of the main text.

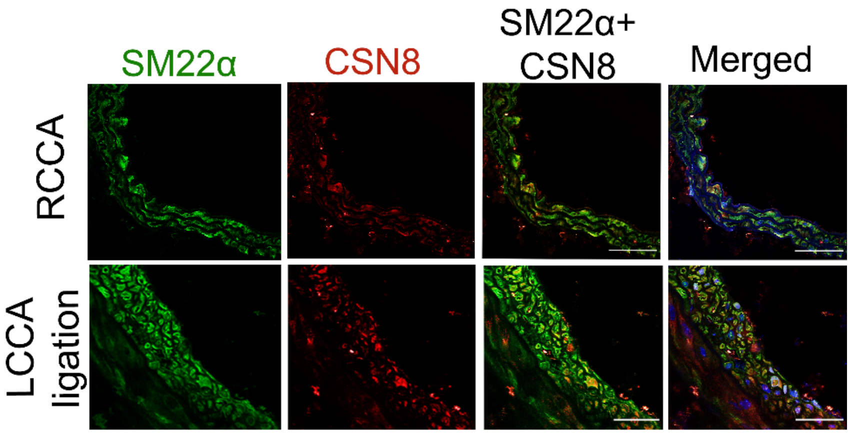


**Figure S2. CSN8 is upregulated in the smooth muscle cells of the left common carotid artery (LCCA) in wildtype mice 1 week after LCCA ligation**. Shown are representative confocal micrographs of LCCA walls immunostained for CSN8 (red) and SM22α (green) as indicated. Nuclei were stained with DAPI (blue). Scale bar= 75μm.

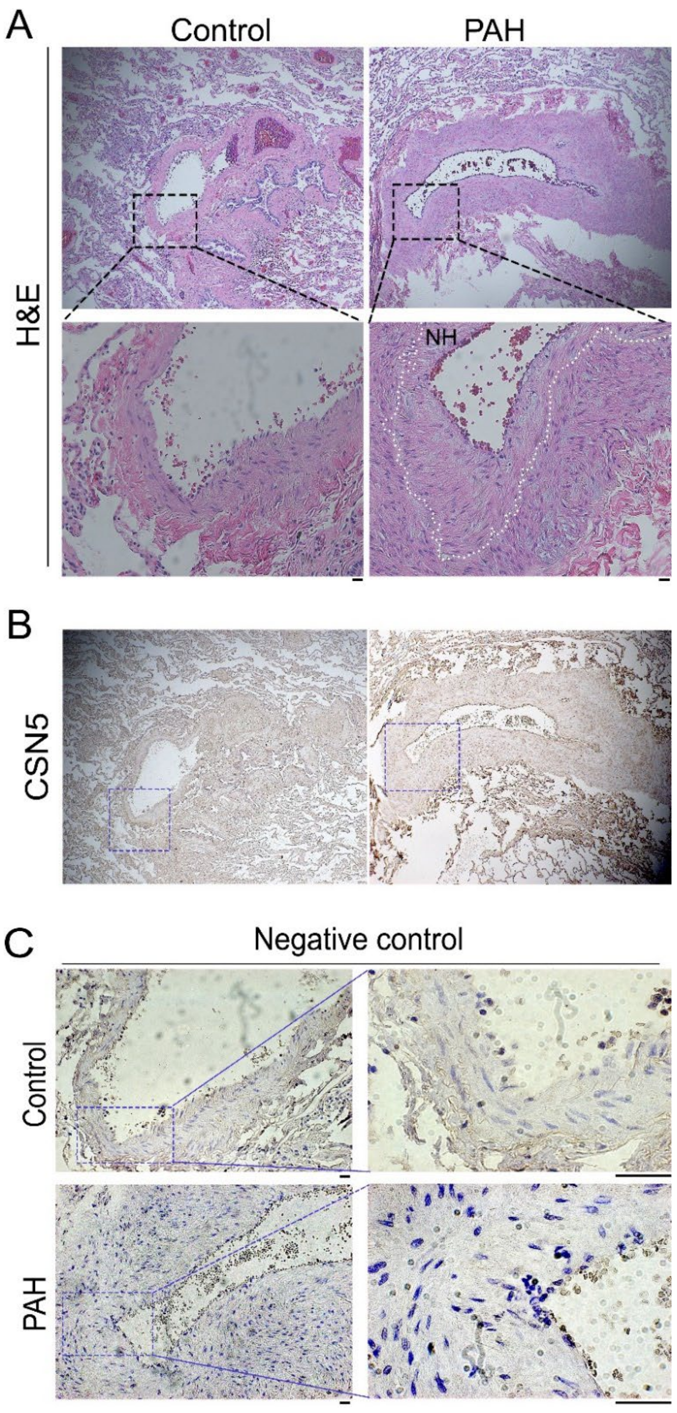


**Figure S3. Expression of CSN5 in human control pulmonary artery (Control) and pulmonary arterial hypertension (PAH) lesions. A,** H&E-stained images of sister sections from the same Control and PAH tissue samples used for the CSN5 immunohistochemistry (IHC) staining as shown in Figure 1F of the main text. **B,** Lower magnification images of the CSN5 IHC stained sections used to collect images shown in **Figure 1F**. **C,** IHC negative control images (for **Figure 1F**) of sister sections with the omission of the anti-CSN5 primary antibody. Scale bar=100μm.

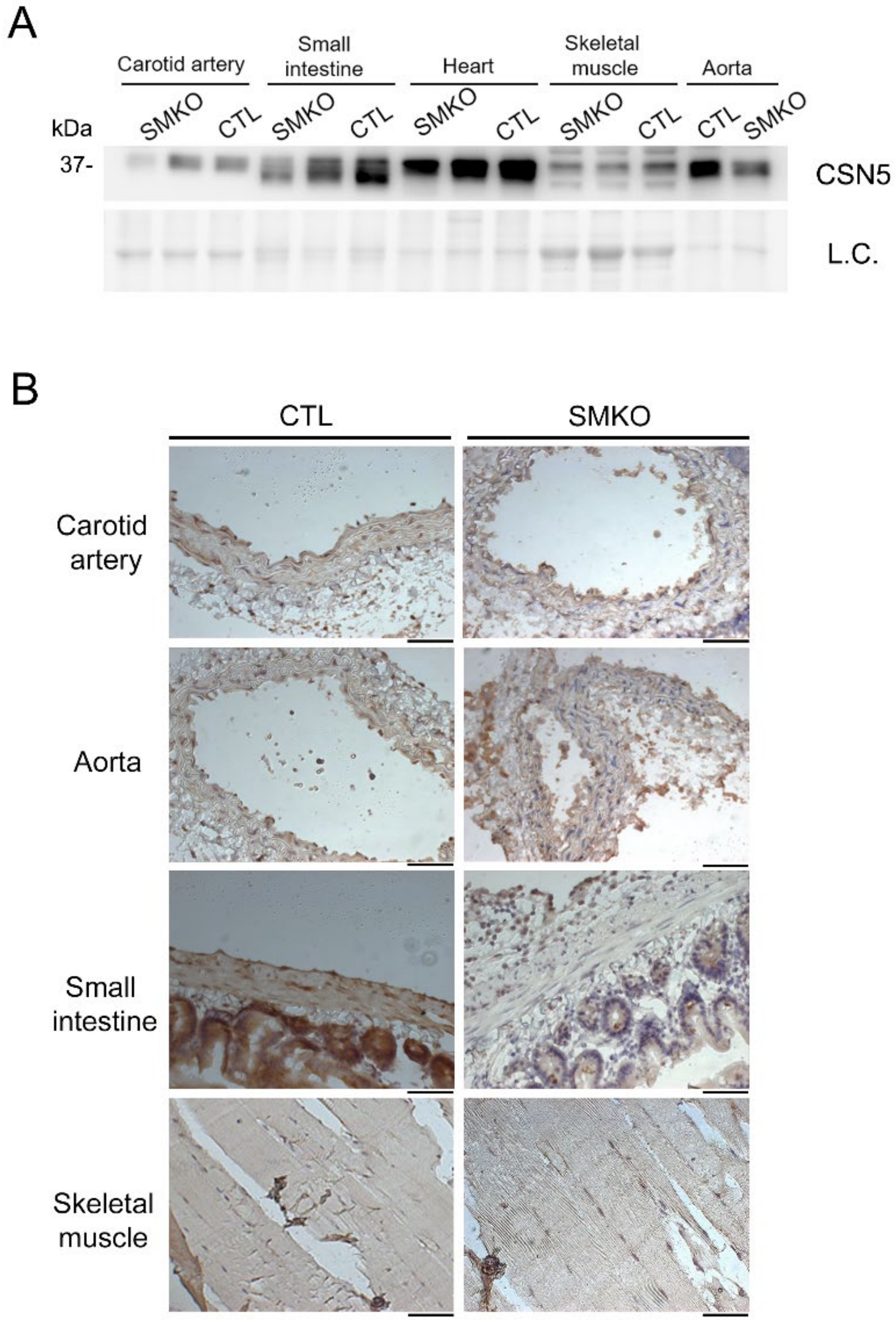


**Figure S4. Characterization of smooth muscle restricted CSN5 knockout (SMKO) in mice.** Two weeks after the last tamoxifen injection, tissues were collected from the homozygous SMKO and the control Myh11-CreERT2 transgenic (CTL) mice. **A** and **B,** Western blot analysis (A) and immunohistochemistry (B) for CSN5 in the indicated smooth muscle-containing organs and non-specific tissues. Scale bar= 100μm.

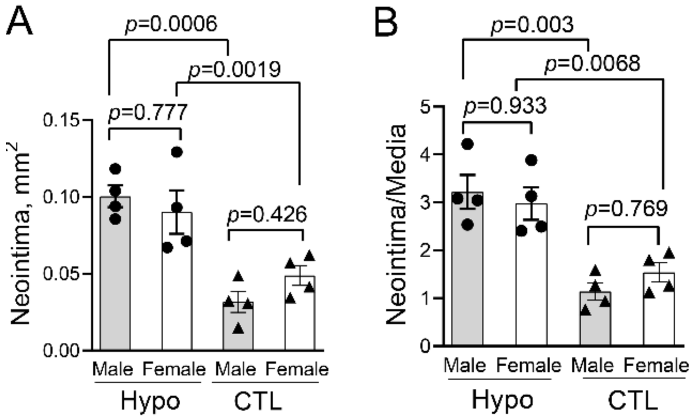


**Figure S5. No sex difference in the exacerbation of LCCA ligation induction of neointimal thickening by CSN8 hypomorphism (CSN8-Hypo) in mice.** LCCAs of CSN8-hypo and CTL mice were collected 4 weeks after LCCA ligation and processed for H&E staining and morphometric analyses as described from Figure 3A of main text. **A** and **B,** Comparison of neointima area (A) and the neointima to media area ratio (B) in the cross-sections of LCCA between male and female mice. Two-way ANOVA shows statistical significance for genotype (CSN8-Hypo vs. CTL) but not for sex (male vs. female). Each dot represents an individual mouse; Mean±SEM.

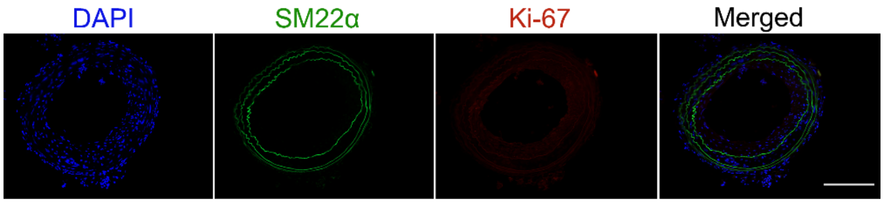


**Figure S6. Confocal micrographs of negative controls for the SM22α and Ki-67 immunofluorescence staining of left common carotid artery (LCCA) sections collected 1 week after LCCA ligation.** The immunofluorescence staining protocol and imaging setting were the same as used for collecting the data shown in Figures 2D, 3H, and 8C of the main text, except for the omission of primary antibodies (SM22α and Ki-67) in (LCCA) after 1 week of ligation. Scale bar=150 μm.

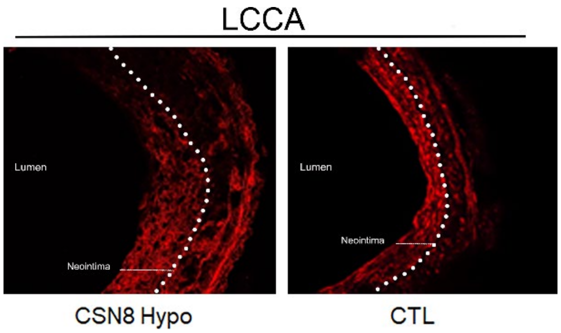


**Figure S7. LCCA ligation-induced neointima formation was exacerbated by CSN8 hypomorphism (CSN8-Hypo) in mice.** One week after LCCA ligation, the ligated arteries from CSN8-hypo mice and littermate controls (CTL) were harvested for analyses. Shown are confocal micrographs immuno-stained for SM22α (red) in the LCCA sections (~2mm proximal to the ligation site).

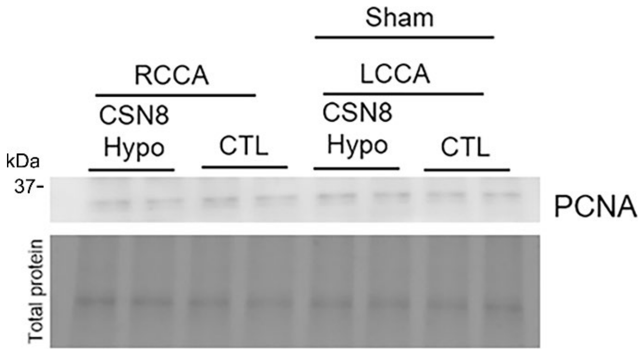


**Figure S8. Representative image of western blot analyses for PCNA in RCCA and LCCA tissues of CSN8-hypo (Hypo) and littermate controls (CTL) mice subjected to LCCA sham surgery.** LCCA and RCCA were collected 1 week after the sham surgery.

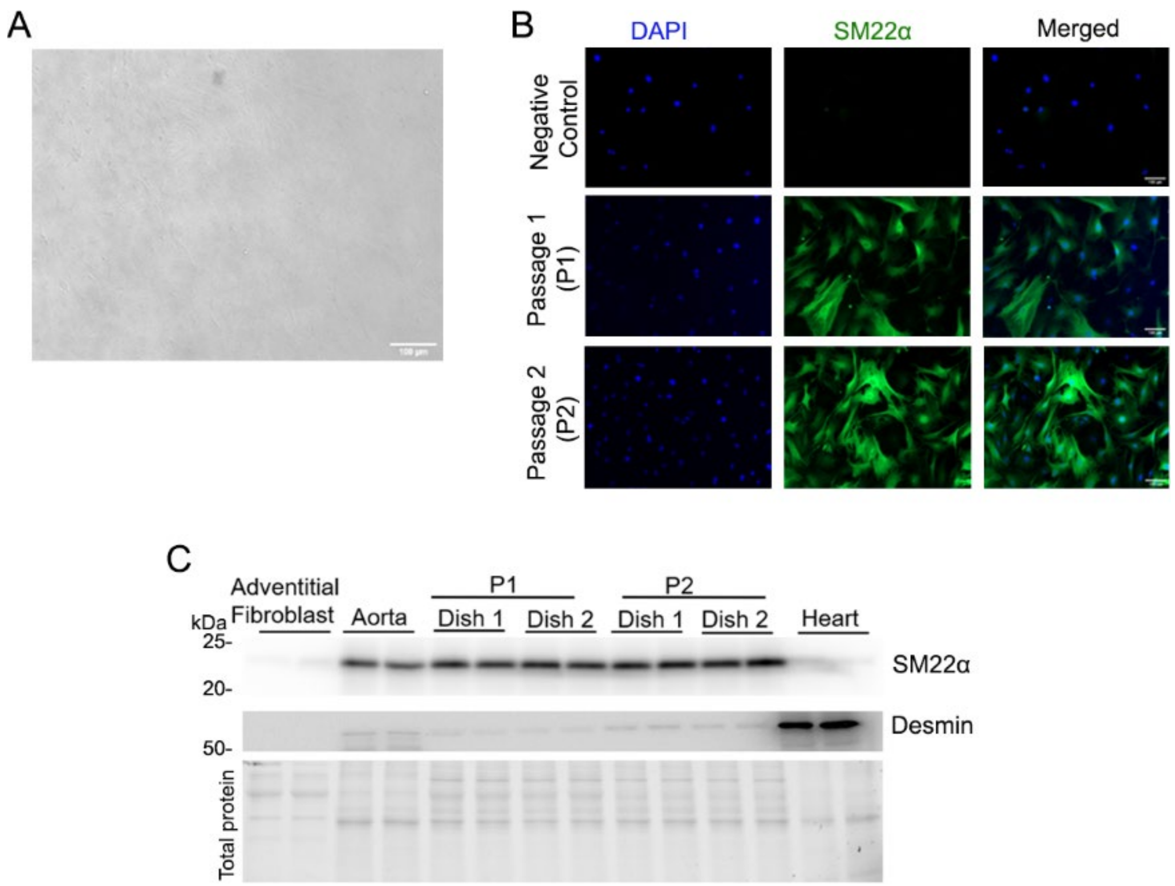


**Figure S9. Validation of cultured vascular smooth muscle cells (VSMCs) isolated from mouse aorta. A**, A representative bright field phase-contrast micrograph of passage 1 (P1) VSMCs in culture (x10)**. B**, Representative micrographs showing the VSMCs in P1 and P2 stained with SM22α (green) and DAPI (blue). **C**, Western blot analysis for SM22α and desmin protein expression in the total protein extracts from cultured adventitial fibroblast, aorta, P1 and P2 VSMCs, and heart tissue**.** Here the heart tissue is used as a positive control for desmin. The lower image in the panel C is the stain-free in-gel labeled total protein image to show the loading.

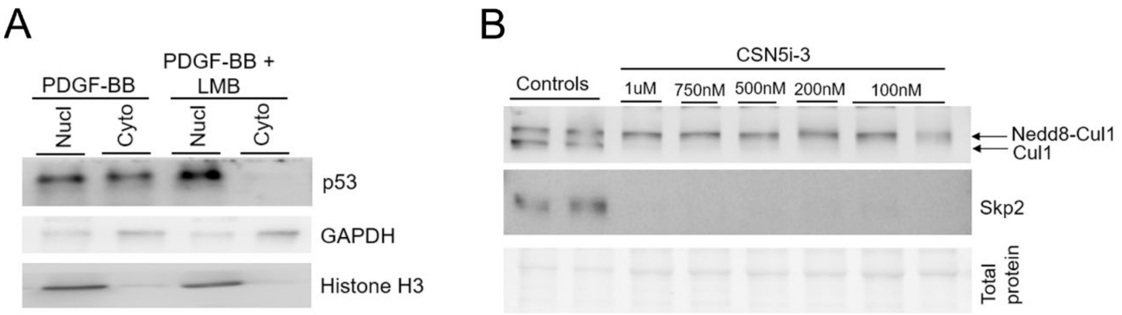


**Figure S10. Verification of the actions of leptomycin (LMB) and CSN5i-3 in cultured VSMCs. A**, Western blot for nuclear and cytosolic p53. VSMCs were treated with PDGF-BB in the absence or presence of LMB (5ng/ml) for 24 h before harvested for extraction of the cytoplasmic (Cyto) and nuclear (Nucl) fractions of proteins. GAPDH and Histone H3 were probed as the markers for cytosolic and nuclear fractions, respectively. **B**, Western blot analyses for neddylated and native Cul1 as well as Skp2. VSMCs in cultures were treated with the indicated concentrations of CSN5i-3 (the small molecule inhibitor of the CSN) for 24 h, then harvested for protein extraction. CSN5i-3 at as low as 100nM completely blocked Cul1 deneddylation and resulted in complete loss of Skp2.

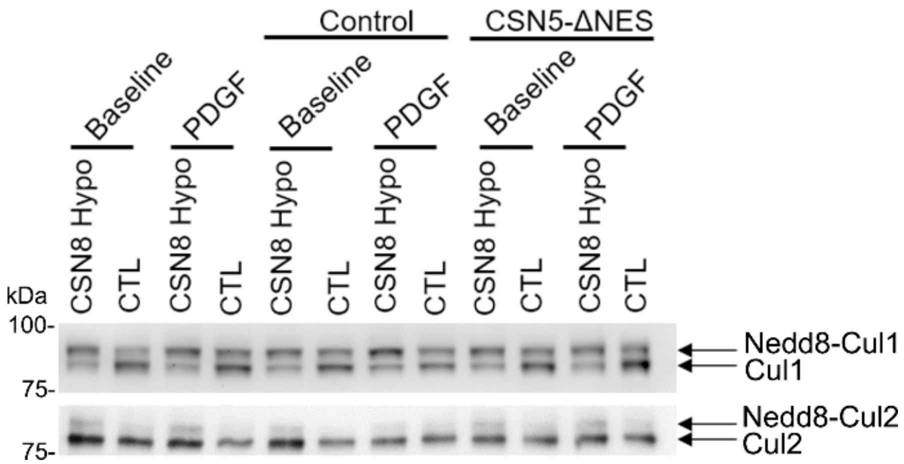


**Figure S11. Western blot analyses for neddylated Cullin 1 (Cul1) and Cul2 proteins in cultured VSMCs transfected with CSN5-ΔNES**. VSMCs were transfected with plasmids expressing EYFP-fused constitutive nuclear CSN5 (CSN5-ΔNES). The plasmid expressing EYFP alone was used as the control. After the transfection, VSMCs were stimulated with PDGF-BB or vehicle control for 24 h before harvested for analyses. Shown are the western blot images probed for Cul1 and Cul2 showing that the transfection of VSMCs with CSN5-ΔNES does not affect the deneddylation function of CSN5.

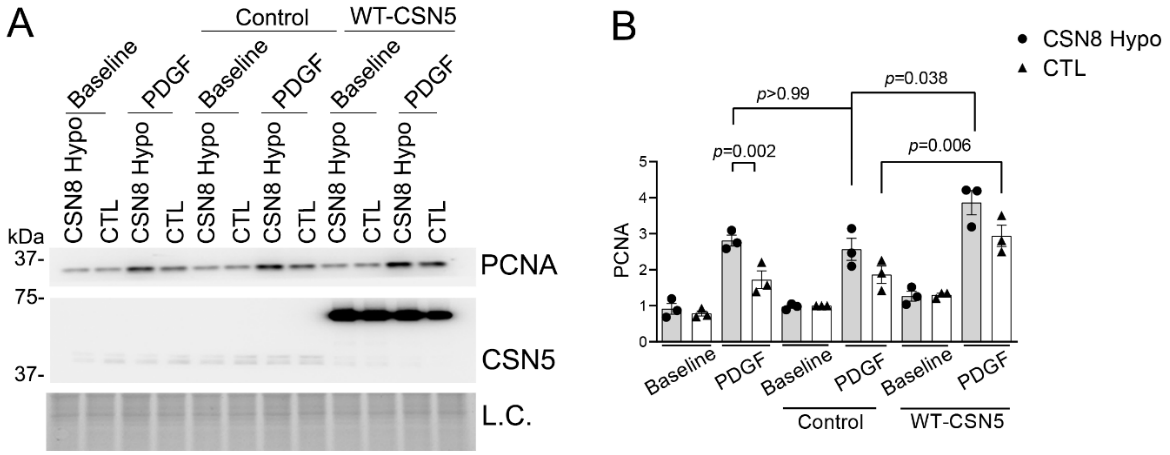


**Figure S12. Overexpression of wild type (WT) CSN5 promotes PDGF-triggered cell proliferation in control and hypomorphic VSMCs.** VSMCs in cultures were transfected with plasmids expressing EYFP fused full-length CSN5 (WT-CSN5) or plasmids expressing EYFP alone (Control). The cells were then treated with PDGF-BB or vehicle control for 24 h and then harvested for protein extraction. **A** and **B**, Representative images (A) of western blot analyses for PCNA and CSN5 and pooled densitometry data of PCNA (B). Mean±SEM; n=3; two-way ANOVA followed by Tukey's tests.
